# Optimal Practice Schedules in a Dual-Rate Model of Motor Adaptation, and Their Recovery by Reinforcement Learning

**DOI:** 10.64898/2026.06.17.732970

**Authors:** Russell Jeter, Dmitrii Todorov, Yaroslav Molkov

## Abstract

A clinician guiding a stroke patient through a 45-minute rehabilitation session, a coach planning a training day, a teacher choosing the order of practice problems, they all face the same question: “given everything practiced so far, what should the next trial be?” The motor-learning literature offers two coarse answers, blocked and interleaved (“random”) practice, with a well-known dissociation, blocked practice gives faster acquisition but worse retention, while interleaved practice gives the opposite. We argue that this dissociation is not a fixed property of practice schedules but a shadow of a richer structure. In particular, for a learner whose memory has a fast shared component and slower context-specific components, the best schedule should be a function of the learner’s current internal state and the time remaining before the retention probe. We make this precise in a minimal two-context fast–slow learner model whose optimal schedules can be computed exactly for short sessions and approximated by a structured beam-search upper bound for longer ones. The optimal schedule is not blocked, not interleaved, and not a single rule; it is a family of schedules determined by how much retention is weighted relative to acquisition. The family has three regimes (alternating, mixed, blocked-with-late-correction) and for long sessions, the optimal schedule has an interpretable structure — exploit one context, repair the neglected one, then interleave to lock in retention. We then investigate whether a reinforcement-learning teacher, observing only the learner’s actions and errors without access to their internal memory states, can learn these optimal policies from interaction alone. Comparing these learned policies against the exact optima, we show that a model-free agent (PPO) recovers the short-horizon schedules and the long-horizon block–repair–interleave motif in the intermediate regime, but the benchmark also exposes a sharp failure in the acquisition-dominated regime, where PPO collapses to pure blocking and misses a sparse terminal correction. A warm-start diagnostic shows this failure is a genuine metastability of policy gradients rather than a tuning artifact, with blocked-plus-switch and pure-blocked acting as competing attractors that PPO cannot stabilize between. A hyperparameter sweep over observation history reveals that the agent requires very little behavioral context to plan optimally, demonstrating that partial observability is not a major barrier to finding optimal practice schedules. Finally, we discuss the implications of our framework for motor adaptation and contextual interference, offering practical insights on how instructors can design finite practice sessions to favor long-term retention.

## 1 Introduction

In motor adaptation experiments, the schedule of practice changes what is learned. The classic finding, replicated across decades and many task families, is the contextual interference effect: blocked practice (one task within a training session) produces faster within-session learning but worse retention than interleaved practice (different tasks each trial), which is harder during the session but more durable afterwards (Schmidt and Bjork, 1992; Guadagnoli and Lee, 2004; Czyż et al., 2024). The same dissociation appears in motor skill training (Shea and Morgan, 1979), in language learning (Nakata and Suzuki, 2019), and in classroom problem sets (Rohrer and Taylor, 2007). The empirical literature is rich, but the theoretical question remains open: when, exactly, should a teacher block, interleave, or do something in between?

This question is not simply academic. A physical therapist working with a stroke patient has a 30-45-minute session, several rehabilitation goals, and an interest in the patient retaining what they practice across days. A coach designing a training schedule has a session, a competition, and a roster of skills. In each case the practitioner is solving a sequencing problem, often guided by intuition: spend a little time on a few items, return to the harder ones, end with a mix. The question we ask is whether this intuition has a principled answer.

### 1.1 Why a Fixed Schedule Cannot Be the Right Answer

One of the classical computational models for motor adaptation is a two-dimensional state space model, where each state corresponds to one of the two motor memory components: a slow-learning and a fast-learning one (Smith et al., 2006). In a learner described by such a model the *fast* process adapts quickly to whatever is being practiced now but decays rapidly between trials, and the *slow* processes adapt slowly but persist (Smith et al., 2006). If practice in different contexts updates a shared fast process and separate slow processes (Lee and Schweighofer, 2009), then the value of practicing context *i* for trial *t* depends not just on the most recent error. It also depends on whether the durable component for context *i* is already saturated (i.e., whether the learner can even still gain from practicing it), and on the alignment of the fast state with the current target. Additionally, it has also been shown that value of practicing specifically depends on the time remaining before the retention is probed in the future (Hadjiosif et al., 2023).

Specifically, we assume a standard experimental paradigm consisting of a training phase of *T* trials, followed by a retention interval (or delay) of Δ trials, and concluding with a retention probe (see, for example, Joiner and Smith, 2008; Lee and Schweighofer, 2009). Although empirical studies organize probe trials in various ways, sometimes omitting them, testing retention immediately, or testing after a delay of hours or days using error clamps or no-feedback trials, their common functional purpose is to measure the durable motor memory that remains after the rapidly-decaying fast component has faded.

To maximize this durable motor memory across multiple contexts, the training schedule must balance both memory processes. Static schedules, however, are poorly suited to this task. A blocked schedule^1^ ignores the slow components of the unpracticed contexts, failing to build durable memory where it is needed. An interleaved schedule, on the other hand, never lets the shared fast state align with any single context long enough to exploit its rapid learning rate. Consequently, a schedule that performs well for such a learner must be state-dependent. The optimal next trial must be chosen as a function of the learner’s current internal state, the fast and slow motor-memory variables (*f, s*_1_, *s*_2_), and not merely of the trial index or the schedule chosen at the outset.

By the learner’s state (equivalently, its motor-memory state or “internal state”) we refer to the vector of fast and slow variables that the state-space model evolves. The teacher, however, never sees these variables. It strictly acts on an externally observed state, a short behavioral history of past trials and outcomes, in the control- or reinforcement-learning sense of a policy. The optimal schedule is state-dependent in the first sense, that is, it is determined by the internal state, whereas the central methodological question of the paper is whether a teacher with access only to the externally observed state can recover it.

The central premise of this paper is the transition from studying static schedules to studying dynamic policies. The right object of inquiry is not a predetermined sequence of trials, but a policy that maps the learner’s current state to the next trial. Once we accept that framing, the question becomes precise: “what does the optimal policy look like, and can a teacher (human or learned) recover it?”

### 1.2 What This Paper Does

We study the simplest setting in which this question has nontrivial structure: a two-context fast– slow learner with one shared fast state and two context-specific slow states. The learning dynamics is simple enough that the optimal practice schedule can be computed exactly for short sessions, by formulating the decision to choose the context to present on a given trial as a mixed-integer optimization problem (Wolsey, 1998) and verifying against brute force (testing all possible context sequences). It is also simple enough that we can obtain a strong heuristic upper bound on the long-session optimum with a structured beam search. Together, the exact solver and the beam search give us ground truth where ground truth is otherwise unavailable in scheduling problems. Solving the schedule exactly is not the goal of this exercise. Rather, the exact optimum provides a benchmark for evaluating the agent’s performance. The unresolved question is whether a teacher, constrained entirely to behavioral observation as in real-world settings, can recover this optimal schedule (or something close).

The paper’s main scientific finding is that the optimal schedules in the learner form a one-parameter family, indexed by *ρ* (the acquisition-retention tradeoff parameter), the relative weight on training error versus retention probe error. As *ρ* grows from small to large, the optimal policy transitions through three qualitatively different regimes:

- **Alternating**, where the optimum is essentially strict alternation;
- **Mixed**, where the optimum is neither alternating nor blocked but a structured combination of both;
- **Blocked with late correction**, where the optimum is blocked practice followed by one or more brief returns to the neglected context.

For long sessions, the family includes a sweet spot in which the optimum has a particularly clean structure consisting of an initial exploitation block in one context, a transition that repairs the neglected slow state, and an interleaved tail right before the retention probe. We call this the block–repair–interleave motif.

The paper’s methodological contribution is that an RL agent, a separate PPO model trained at each *ρ* which observes only the learner’s behavior and never its internal state, recovers the regime structure that the exact solver produces, including the block–repair–interleave motif for long horizons. A history-depth hyperparameter sweep also shows that the agent does not need privileged access to the latent fast and slow states; a small window of past actions and errors is sufficient to learn these schedule motifs. We train a separate specialist agent for each objective (*T, ρ*) rather than a single generalist. This isolates the recoverability question and is a deliberate scope limit, since a real clinician would not retrain per objective (Section 7.3).

### 1.3 Contribution

The learner is deliberately minimal. We do not claim that the schedules optimal in it are the schedules optimal in a stroke patient or a baseball pitcher. The benchmark’s role is to expose the shape of the scheduling problem including the existence of a state-dependent optimal policy, the regime structure across the acquisition-retention tradeoff parameter *ρ*, and the question of whether an agent can recover this structure with realistic observability. The benchmark’s simplicity allows for an exact solver to compute the optimal schedules. Without explicit knowledge of the optimal schedules, every claim about RL would either be unfalsifiable (if there were no ground truth) or under-motivated (if the only baseline were another empirical agent). The “Related Work” section contextualizes our contribution with the most directly relevant prior literatures.

### 1.4 Roadmap

Section 2 reviews the motor-learning, optimal-control, machine-teaching, and partial-observability literatures the paper draws on. Section 3 states the phenomenon and the question more precisely. Section 4 introduces the learner. Section 5 describes the exact solver and the beam-search bound. Section 6 characterizes the optimal-schedule manifold and the block–repair–interleave motif. Section 7 sets up the reinforcement-learning agent. Section 8 reports what the agent recovers at short horizons; Section 9 contains the long-horizon comparison and the schedule-strip evidence for the block–repair–interleave motif; Section 10 characterizes the observability the agent requires. Section 11 documents the failure modes the exact solver makes visible. Section 12 returns to motor adaptation, contextual interference, and the practical implications for finite-time practice scheduling.

## 2 Related Work

The paper draws on four loosely connected literatures: dual-rate models of motor adaptation, contextual-interference and variability-of-practice studies, optimal-control and machine-teaching formulations of training schedules, and reinforcement learning under partial observability. We summarize each only as deeply as our specific claims require.

### 2.1 Dual-Rate Motor Adaptation

The fast–slow memory architecture our benchmark instantiates was introduced by Smith et al. (2006) as the two-state model of force-field adaptation, capturing savings, spontaneous recovery, and the dissociation between rapid acquisition and durable retention. In their framework, spontaneous recovery is demonstrated when a brief opposite perturbation extinguishes net adaptation, but a subsequent error-clamp reveals the persistent slow state. Similarly, a short re-exposure to the original perturbation triggers a rapid recovery of performance because the slow process retains the memory of the initial adaptation. Lee and Schweighofer (2009) extended the framework to multi-context settings with a parallel architecture — shared fast learning and context-specific slow learning — which is exactly the architecture our learner uses. Berniker and Kording (2011) formalized the inference problem behind multi-state adaptation, and Heald et al. (2021, 2023) subsequently recast it in a contextual-inference framework. The learner is the smallest two-context instance of this lineage.

### 2.2 Contextual Interference and Variability of Practice

The contextual-interference (CI) effect was named by Shea and Morgan (1979) and recast as a general schedule-design principle by Schmidt and Bjork (1992). The challenge-point framework (Guadagnoli and Lee, 2004) formalized the tradeoff between immediate task difficulty and durable learning gain. The broader variability-of-practice tradition (Wulf and Shea, 2002) cautioned that principles derived from simple-skill experiments do not always generalize to complex skill learning. A recent meta-analysis (Czyż et al., 2024) confirms that high-CI practice improves retention across motor tasks. Our framework reframes the CI dissociation as a boundary case of state-dependent control (a principle prominent in architectures like MOSAIC (Haruno et al., 2001)). Blocked and interleaved are the endpoint regimes of a *ρ*-indexed manifold whose interior is rarely tested experimentally because conventional designs hold the schedule family fixed.

### 2.3 Optimal Scheduling for Motor-Learning Models

Optimal scheduling in motor learning has evolved from performance-based heuristics (e.g., Choi et al., 2008) to formal optimal control treatments. The closest precursor to our work is by Lee et al. (2016), who applied Pontryagin’s maximum principle to a continuous relaxation of the binary scheduling decision under dual-rate dynamics. However, differences in both the objective and task structure distinguish our findings. First, while Lee et al. (2016) optimize solely for final retention, our framework incorporates a trial-by-trial training error penalty, parameterized by the acquisition-retention tradeoff parameter *ρ*. Second, they study asymmetric task difficulties and find that optimal schedules distribute trials of the harder task evenly throughout the session. In contrast, we study symmetric tasks under a joint objective, showing that the optimal schedule transitions through discrete regimes—including the block–repair–interleave motif, which balances early training performance with late retention. Finally, while their optimal control law requires full observability of the learner’s internal state, our reinforcement learning teacher operates under partial observability, relying only on the trial history of past actions and errors.

### 2.4 Machine Teaching and Adaptive Tutoring

Treating the teacher as a sequential decision-maker that optimizes a student outcome has a substantial recent literature. Machine teaching (Zhu, 2015) frames teaching as the inverse problem to machine learning: choose an optimal training set given a known student. Curriculum learning for RL (Bengio et al., 2009; Narvekar et al., 2020) studies orderings that accelerate downstream training. RL-based adaptive tutoring (Bassen et al., 2020; Reddy et al., 2017) deploys learned policies to schedule educational tasks, typically without a closed-form optimum to evaluate against. Closer to the active-learning side, Peltola et al. (2019) formalize machine teaching of active sequential learners. Our learner contributes a benchmark in this space with a computable optimum, which is necessary to evaluate the performance of a learned policy against the absolute theoretical limit.

### 2.5 Spaced Repetition and Forgetting-Aware Scheduling

A parallel line of research in educational technology schedules practice to optimize delayed retention against explicit models of forgetting. For instance, Pavlik Jr and Anderson (2008) utilized ACT-R memory dynamics to derive optimal practice intervals, while Settles and Meeder (2016) fit half-life regression models to language-learning data to schedule review sessions. While these approaches focus on memory decay over time rather than multi-context fast-slow interaction, they share our core optimization objective of maximizing performance at a delayed retention test. This corresponds directly to the retention-dominated (small-*ρ*) regime of our framework.

### 2.6 Partial Observability and History-Based RL

Our teacher observes a behavioral history of past actions and errors but not the latent fast–slow state, which formally makes the scheduling problem a partially observable Markov decision process (Kaelbling et al., 1998). The history-window observation (10) is one classical solution; recurrent policies (Hausknecht and Stone, 2015) are another. We use the windowed-MLP version because it provides a natural hyperparameter sweep over the error history length and context history length (*K, M*), that lets us measure how much behavioral history is enough; the recurrent variant is left for follow-up work. The framework’s affinity with POMDP control is part of what makes it a clean RL-methodology benchmark.

## 3 The Phenomenon and the Question

Consider a single practice session of *T* trials with two contexts. After the session, the learner is evaluated for retention across both contexts after a delay Δ. The session’s quality is summarized by a weighted sum:

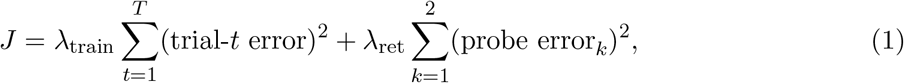

where *λ*_train_ weights performance during the session and *λ*_ret_ weights probe error. The probe error in context *k* is the residual that remains when, after the Δ-trial retention interval, the learner is tested once on context *k*. We assume that no learning occurs during this interval or at the probe. Across the Δ delay trials the memory states evolve by decay alone, with no error update, and each probe is a single no-feedback trial that reads out the learner’s response without updating any state. The two probes therefore share the same post-delay memory state, and the probe error does not depend on the order in which the contexts are tested. The parameter that ultimately controls the structure of the optimal schedule is the acquisition-retention tradeoff parameter, defined as the ratio

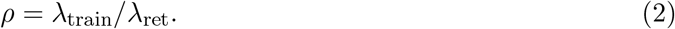

At small *ρ* the teacher cares mostly about retention; at large *ρ* the teacher cares mostly about within-session performance. Real learning settings span the range of goals. Because the objective sums *T* training-error terms against only two probe terms, the two contributions carry equal total weight near *ρ* ≈ 2*/T* (about 0.1 at *T* = 20 and 0.01 at *T* = 200), not at *ρ* = 1. A clinical session in which only post-session function matters has *ρ* near zero; a session that cares chiefly about within-session performance has *ρ* ≫ 2*/T*. The schedule transitions studied below all fall in *ρ* ∈ [0.01, 0.07] at *T* = 20, consistent with this scaling.

This objective function motivates two primary questions:

1. For any given value of the acquisition-retention tradeoff parameter *ρ*, what is the practice schedule that minimizes the joint cost *J* ?
2. How does the optimal schedule transition between blocked, interleaved, or other distinct sequencing regimes as a function of *ρ*?

The standard schedule families (blocked, alternating, random) are useful empirical categories, but they cannot answer either question if the optimal schedule depends on the learner’s state.

### 3.1 Why a State-Dependent Answer Is Plausible

A simple thought experiment makes the case that a state-dependent schedule is plausible. Suppose the learner has just spent 30 trials practicing context 1. The fast component is well-aligned with context 1; the slow component for context 1 is also doing most of the work. Context 2 has not been practiced, so its slow component is still at zero. No durable memory has been built for its slow component, and it carries its full durable gap.

#### 3.1.1 “What should the next trial be?”

If *ρ* is large (training-error-dominated), the answer is context 1 again: switching costs immediate performance, and the retention probe is far enough away that retention can be repaired later. If *ρ* is small (retention-error-dominated), the answer is context 2: the durable gap is bigger there, and a single trial of context 2 can repair it cheaply because the slow process is sensitive but slow. At intermediate *ρ*, the answer depends on details of the state.

This is the structure we make precise. We use the smallest model that exhibits it.

## 4 A Minimal Model of Fast–Slow Memory

We consider the case of a fast–slow learner in 1D flat continuous space with two contexts (*K* = 2) corresponding to two opposing perturbations of equal magnitude, *p*_1_ = +1 and *p*_2_ = −1, that the learner must compensate for. It is a 1F2S model in the terminology from Lee et al. (2016). The motor-adaptation reading is two opposing force fields, or clockwise and counterclockwise visuomotor rotations, applied to reaches toward a common kinematic target. It is not necessary for these perturbations to be strictly opposing, but this stark contrast is the worst possible case for learning in two contexts. For simplicity, we assume that the learner receives feedback about its action only after an action is performed (not during it). In motor adaptation experiments it is often implemented using endpoint-only feedback or shooting/throwing movements (Krakauer et al., 2019).

Without loss of generality we take both contexts to reach the same target located at 0 (a recentering of coordinates). We further assume, as a modeling simplification following Lee and Schweighofer (2009), that the fast process generalizes perfectly between the contexts while the slow states do not generalize at all. We employ the standard state space model and describe it mathematically below.

First, the learner’s internal (motor-memory) state is described as a three-dimensional vector

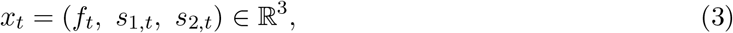

where *f*_*t*_ is a single shared fast state and *s*_1,*t*_, *s*_2,*t*_ are two context-specific slow states. When context *i* is practiced during trial *t*, the learner produces an adaptive motor command, whose output endpoint has the coordinate

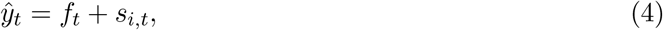

and incurs the residual error *e*_*t*_ = *p*_*i*_ − *ŷ*_*t*_. This continues for *T* trials. On the first trial of a never-practiced context this residual is the full perturbation magnitude (*e*_*t*_ = *p*_*i*_); it shrinks as *f*_*t*_ + *s*_*i,t*_ approaches *p*_*i*_ over subsequent trials. The internal states update by

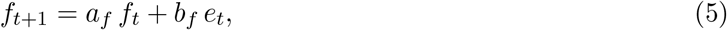

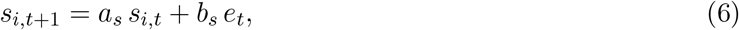

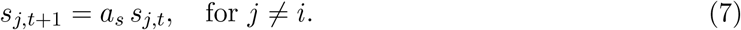

Here, the retention parameters (*a*_*f*_, *a*_*s*_) describe memory retention between trials, and the learning rates (*b*_*f*_, *b*_*s*_) describe sensitivity to error (adaptation rate). The four parameters (*a*_*f*_, *b*_*f*_, *a*_*s*_, *b*_*s*_) satisfy 0 *< a*_*f*_ *< a*_*s*_ *<* 1 and *b*_*f*_ *> b*_*s*_ *>* 0, such that the fast state learns strongly but decays rapidly while each slow state learns weakly but persists. We use the following default parameter values, along with a retention interval (or delay) of Δ = 3 trials before the retention probe:

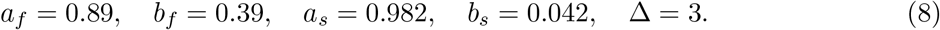

These specific values are chosen to expose the rich, non-trivial transitions of the optimal schedules. In particular, they ensure a wide range of the tradeoff parameter *ρ* where the optimal schedule is neither pure blocking nor strict alternation, but rather a structured, state-dependent mixture of both (characterized in Section 6).

### 4.1 Why These Dynamics, and Where They Come From

The fast–slow split is the structural ingredient identified by the dual-rate motor-learning literature (Smith et al., 2006) as load-bearing for many adaptation phenomena. The shared-fast/context-specific-slow architecture, which gives the learner its two-context structure, is the form that empirical work on dual adaptation and contextual interference converges on (Lee and Schweighofer, 2009; Lee et al., 2016). The learner model presented here compresses this lineage into the smallest possible setting. It has enough structure to make the scheduling problem nontrivial but not so much that the optimal schedule is hidden by behavioral noise or by an over-parameterized state space.

The minimality is a feature, not a limitation. Our claim is not that the learner faithfully captures any particular real adapter; it is that it is the smallest setting in which the question “when should a teacher switch?” admits a precise, computable answer. If the answer is interesting here, that is a reason to ask the same question in more dynamically rich models. If the answer were trivial, that is, if the optimum were always blocked or interleaved, the question would be settled for this model, and a more complicated model would be necessary to exhibit state-dependent optimal learning schedules.

### 4.2 Why the Model Can Be Solved Exactly

Two properties of the learner make the optimal schedule computable. The dynamics are linear conditional on the action sequence, so the trajectory of (*f*_*t*_, *s*_1,*t*_, *s*_2,*t*_) is a deterministic function of the schedule. The action space is binary (at any trial the policy chooses between two contexts to present), so the schedule space has size 2^*T*^. For modest *T* this is small enough to enumerate by brute force (*T* ≤ 22 for our hardware) and tractable for a mixed-integer reformulation up to *T* ≈ 32. Beyond that, the search space is too large to guarantee a minimum in our environment, and we use a structured beam search as a tight heuristic upper bound. Section 5 covers the details for readers who want them.

## 5 Computing the Optimal Schedule

For short sessions, we solve the binary scheduling problem by mixed-integer reformulation (Wolsey, 1998). Let *c*_*t*_ ∈ {0, 1} denote “at trial *t* we present context 1”. The error can then be computed using the following equation

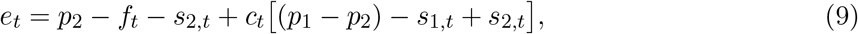

and the trajectory (*f*_*t*_, *s*_1,*t*_, *s*_2,*t*_)_1:*T*_ is a deterministic function of *c*_1:*T*_. Because the learner is deterministic with a fixed initial condition, the optimal open-loop (no knowledge of the model’s state) sequence and the trajectory of the optimal closed-loop state-feedback policy coincide. We therefore compute optimal schedules directly and reserve the word “policy” for the conceptual closed-loop problem and the RL teacher. For *T* ≤ 22, exhaustive enumeration of all 2^*T*^ schedules is feasible and forms the basis used to produce every figure about exact results in this paper.

For sessions of *T* = 200 trials, the search space contains 2^200^ schedules, causing the mixed-integer programming formulation to stall. To overcome this, we apply a beam search seeded with structured candidate suffixes (alternation, blocking, single-switch, and repeated greedy last-trial rollouts) and subsequently refined via single-flip and switch-boundary moves. The result is a strong heuristic upper bound rather than a certified optimum; we report it as such throughout. Appendix A provides the beam hyperparameters and a search space sensitivity analysis showing that the overall macrostructure survives the removal of any single suffix family.

## 6 The Optimal-Schedule Manifold

The exact solver and the beam search produce more than point optima; they trace a *ρ*-indexed manifold of optimal (and, at long horizons, near-optimal) schedules. This manifold is the central scientific contribution of this paper.

### 6.1 Three Regimes for Short Sessions

For sessions of *T* = 20 trials, the optimal schedule transitions through three distinct regimes as *ρ* increases from 0.01 to 0.07 (Figure 1); Table 1 summarizes them.

**Table 1:**
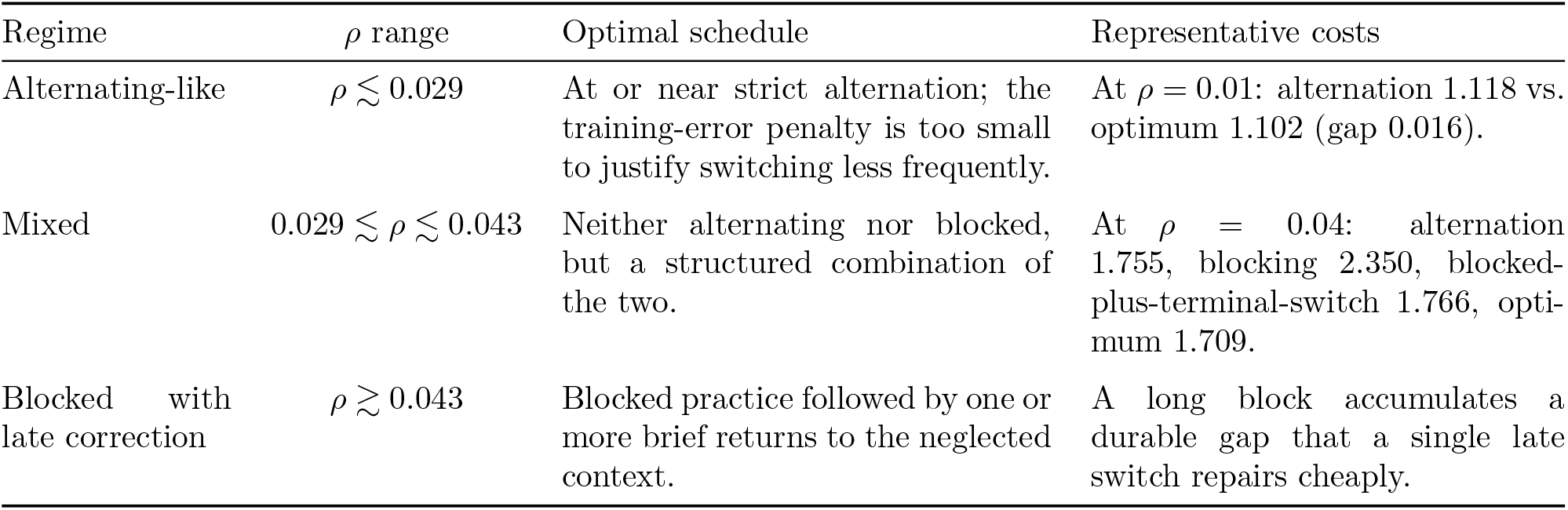
The three optimal-schedule regimes at *T* = 20, as the acquisition-retention tradeoff parameter *ρ* increases from retention-dominated to acquisition-dominated. Costs are the total objective *J*; “optimum” denotes the exact brute-force minimum. The mixed regime is the diagnostic one. In it, no element of the static endpoint family (alternating, blocked, or blocked-plus-terminal-switch) attains the optimum.

| Regime | $\rho$ range | Optimal schedule | Representative costs |
| --- | --- | --- | --- |
| Alternating-like | $\rho \lesssim 0.029$ | At or near strict alternation; the training-error penalty is too small to justify switching less frequently. | At $\rho = 0.01$: alternation 1.118 vs. optimum 1.102 (gap 0.016). |
| Mixed | $0.029 \lesssim \rho \lesssim 0.043$ | Neither alternating nor blocked, but a structured combination of the two. | At $\rho = 0.04$: alternation 1.755, blocking 2.350, blocked-plus-terminal-switch 1.766, optimum 1.709. |
| Blocked with late correction | $\rho \gtrsim 0.043$ | Blocked practice followed by one or more brief returns to the neglected context. | A long block accumulates a durable gap that a single late switch repairs cheaply. |

**Figure 1:**
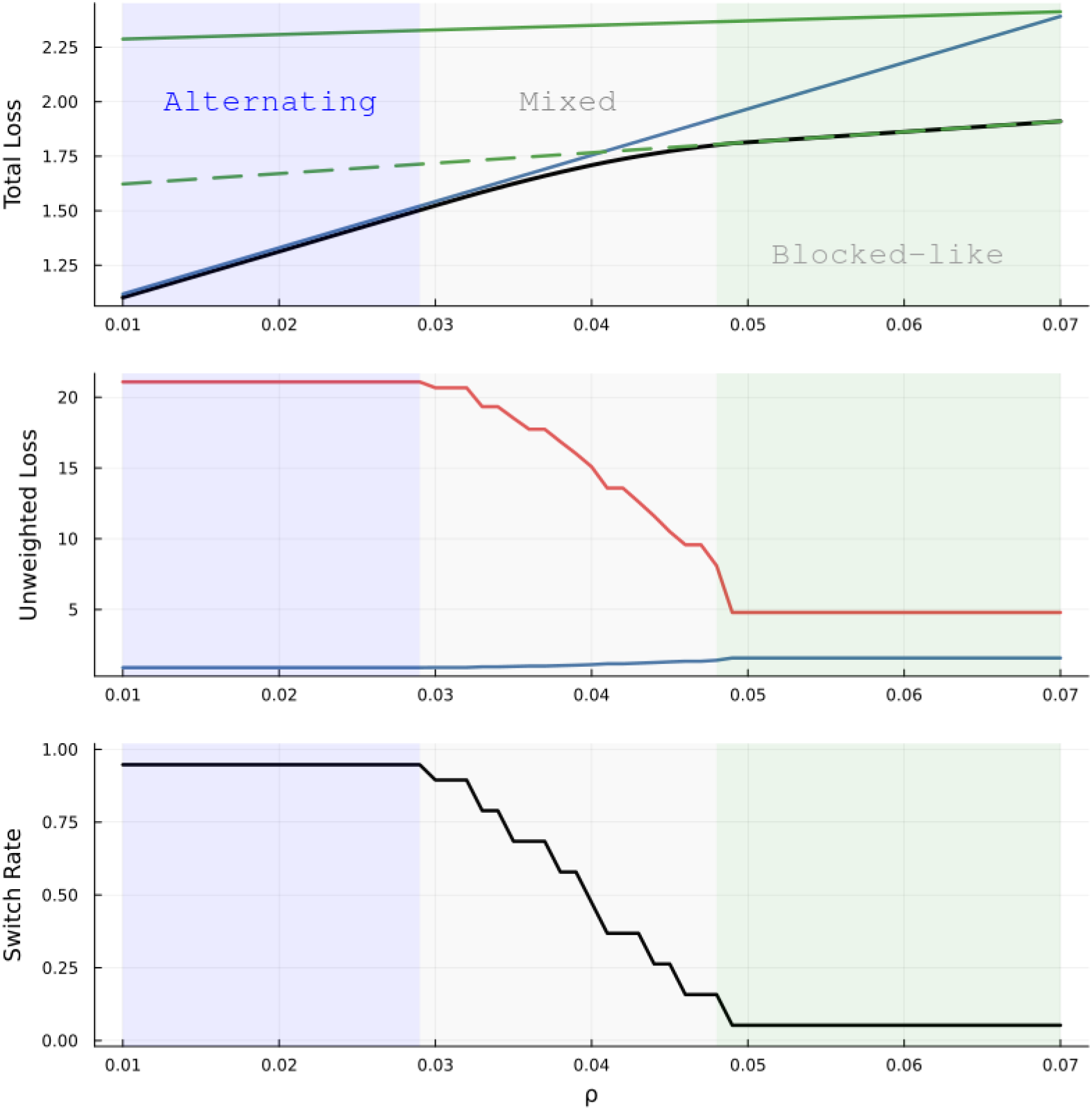
Exact search results over the parameter *ρ* for a short session (*T* = 20 trials). The top panel shows the total loss of the exact optimum (solid black) compared to static baselines: alternating (solid blue), blocked (solid green), and blocked plus a single terminal switch (dashed green). The middle panel decomposes the exact optimum’s total loss into its two unweighted components: the training loss (red) and the probe/retention loss (blue). The bottom panel shows the switch rate of the exact optimum as a function of *ρ*, transitioning from alternating-like behavior at small *ρ*, through a mixed transitional phase in the interior, to blocked practice at large *ρ*. The mixed-optimum interior cannot be matched by any element of the static endpoint family.

The transition is progressive in *ρ*, monotone in the switch rate. The schedule’s switch rate, the percent of trials that a different context from the previous is selected, decreases monotonically from about 0.947 (18*/*19) in the alternating regime to near 0 in the blocked regime, with the mixed interval serving as a complex transitional phase. Even in the retention-dominated limit the optimum is not exactly strict alternation. Because the shared fast state has only partially decayed by the probe (a factor 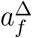), it is read into both probe errors with the same sign, whereas the two contexts have opposing targets; the optimum therefore arranges the final one or two trials to balance this residual, a near-alternation with a terminal correction that beats strict alternation by about 0.015 at *ρ* = 0.01 (switch rate 0.947 rather than 1.0). As the retention delay Δ grows and the fast state washes out before the probe, this correction vanishes and strict alternation becomes exactly optimal. Figure 2 shows the optimal schedule in this mixed regime (*ρ* = 0.04), which is neither strict alternation nor pure blocking.

**Figure 2:**
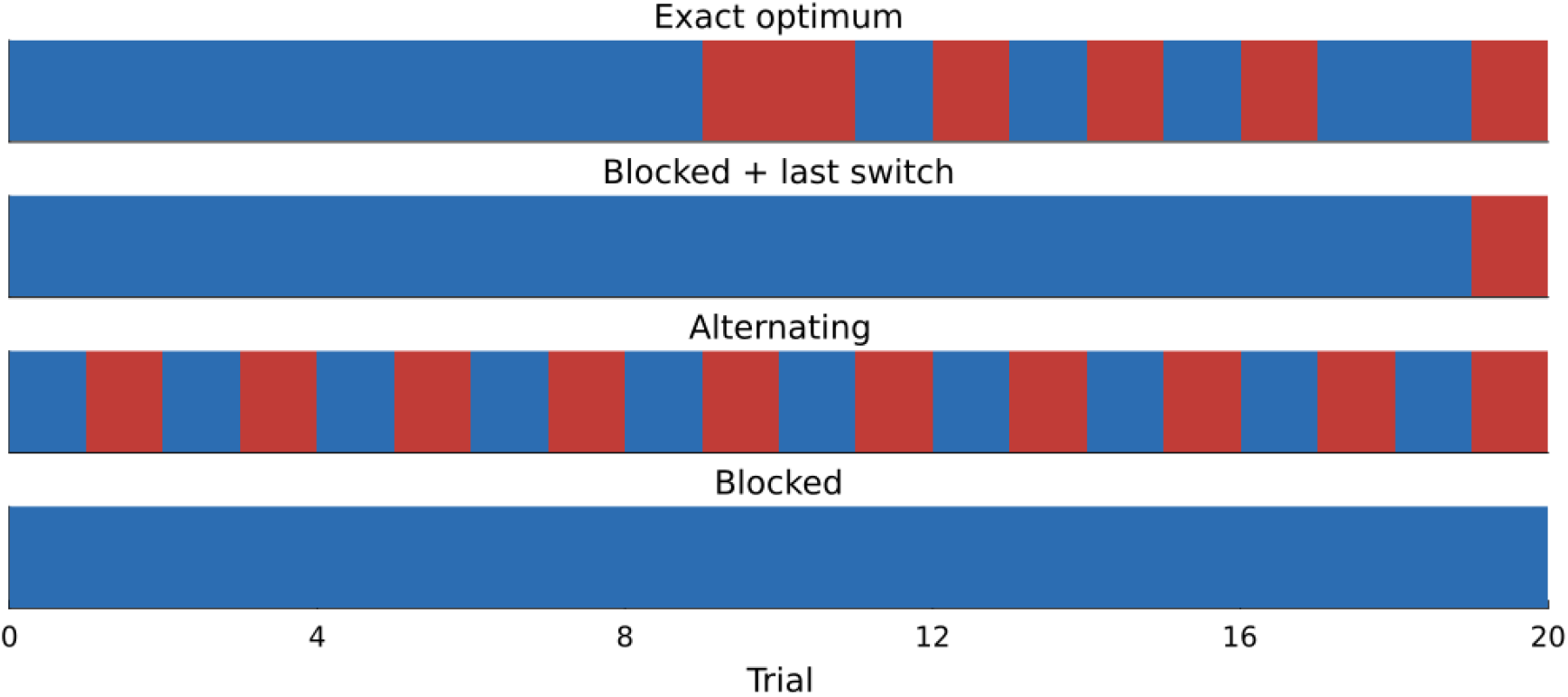
Optimal schedule structure in the mixed regime (*T* = 20 trials, *ρ* = 0.04). The exact optimum is a state-dependent sequence that is neither strict alternation nor pure blocking. A blocked schedule with a single terminal switch (blocked-plus-switch) captures the late-repair mechanism near the probe, but fails to match the detailed structure of the global optimum.

The phase structure is preserved as the horizon grows: longer sessions admit alternating-like optima at larger *ρ* and blocked-with-late-correction optima at larger *ρ* still, with the mixed regime widening (Figure 3). The qualitative picture, however, is the same: as the teacher’s focus moves from retention-dominated to acquisition-dominated, the optimal schedule moves progressively from alternating through mixed to blocked-with-correction.

**Figure 3:**
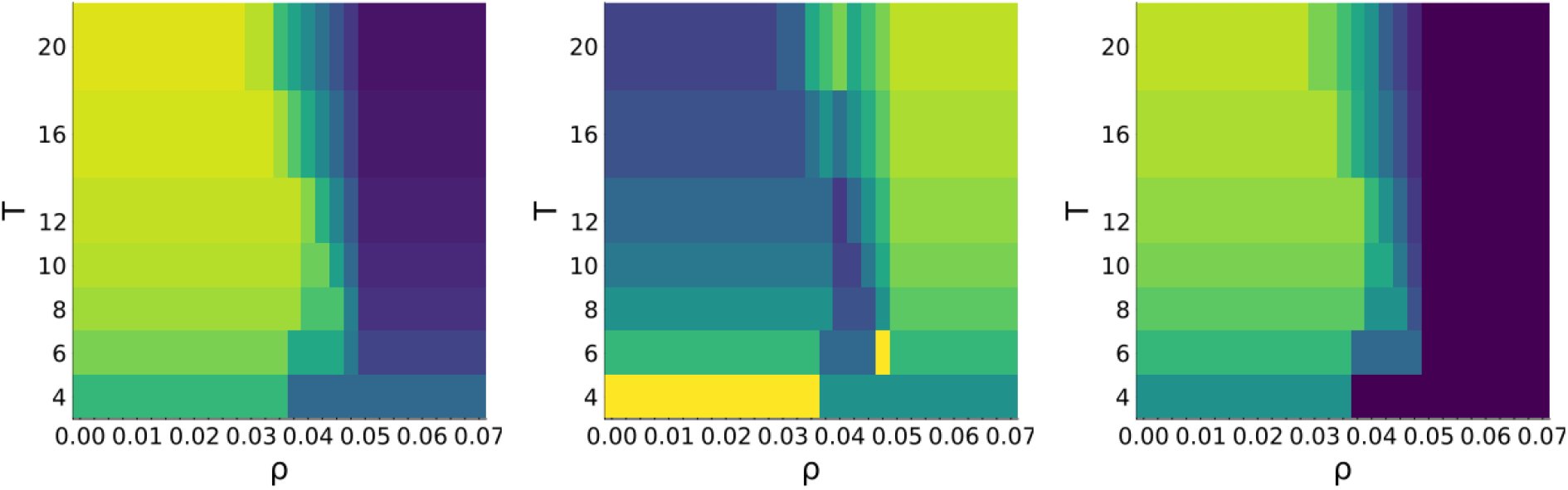
The horizon–*ρ* phase diagram of optimal schedules, showing the switch rate (left), the Hamming distance to alternating schedules (middle), and the Hamming distance to the blocked-plus-switch baseline (right). In all panels, yellow corresponds to larger values (high switch rate or large Hamming distance) and dark purple to smaller values (low switch rate or small Hamming distance near zero). In the left panel, yellow indicates strict alternation; in the middle panel, dark purple indicates schedules matching the alternating baseline; and in the right panel, dark purple indicates schedules matching the blocked-plus-switch baseline. Schedule class is computed by Hamming distance to alternating, blocked, and blocked-plus-switch endpoints; “mixed” indicates schedules outside the endpoint family. The three-regime structure is preserved across horizons, with longer sessions widening the mixed interval.

### 6.2 The Block–Repair–Interleave Motif

For long sessions, the optimal schedule exhibits a particularly clean structure. Figure 4 illustrates the best beam-search schedule alongside two diverse near-optimal alternatives at (*T, ρ*) = (200, 0.013), representing an ideal long-horizon regime. All three share the same distinct macrostructure: a long initial exploitation block in one context, a corrective transition to the other, and an interleaved tail near the probe. We interpret this sequence as follows:

**Figure 4:**
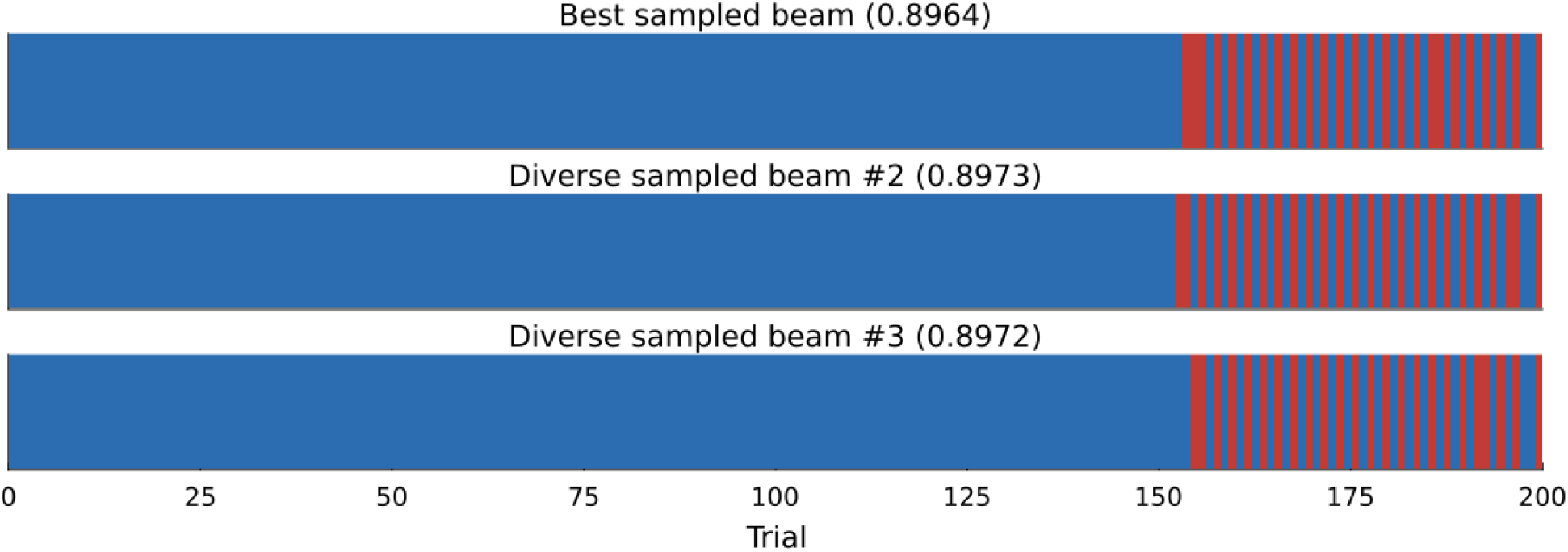
The block–repair–interleave motif at (*T, ρ*) = (200, 0.013). The best beam schedule and two diverse near-optimal alternatives all share an initial exploitation block, a corrective transition, and an interleaved tail. The interpretation is mechanical: exploit the fast state when it is willing, repair the neglected slow state when the cost is low, and stabilize both contexts near the probe.

1. *Block*. At the start of the session, the fast state is uncommitted and both slow states are zero. Repeatedly practicing a single context is the most efficient way to drive its prediction toward the target.
2. *Repair*. After approximately 100 trials, the fast state and the active slow state become saturated; further trials in that context accrue training error without yielding significant retention gains. Meanwhile, the unpracticed context still has no durable memory: its slow state remains at zero, so it carries its full durable gap. A brief sequence of trials in the neglected context, executed while the fast state remains highly responsive, effectively repairs this durable gap.
3. *Interleave*. During the final ≈ 50 trials, once the durable component is approximately corrected for both contexts, alternating practice maintains both near the probe target while minimizing training cost.

This motif persists throughout our search-space sensitivity analyses. Removing any single suffix family from the candidate pool or introducing novelty pressure alters the fine-grained details of the schedule, but leaves the overall qualitative structure intact. We detail these sensitivity analyses in Appendix A to confirm that the observed motif is fundamentally driven by the objective function itself, rather than being an artifact of a particular search heuristic.

The phase structure of Figures 1–4 is independent of the agents we train later; it is a direct consequence of the fast–slow learner’s dynamics. The structure of these optimal and near-optimal schedules implies that for any learner of this general type, the choice between blocked and interleaved practice has no fixed answer. The right choice is a continuous function of *ρ*, spanning three qualitatively different regimes. In this framework, the contextual interference effect is not a static property of practice schedules, but a shadow of the underlying optimal control problem of balancing fast-acting acquisition against slow-decaying retention. We turn next to whether a learning agent can recover this structure from interaction alone.

## 7 The Reinforcement Learning Setup

We treat the teacher as a reinforcement-learning (RL) agent acting on the learner. At each trial the agent picks an action *a*_*t*_ ∈ {1, 2} corresponding to motor perturbations, the learner updates, and the agent receives the negative trial error squared as reward; at the end of the session the agent receives the additional retention-error reward. The agent’s total return is −*J* (*a*_1:*T*_), the same objective the exact solver minimizes.

### 7.1 What the Agent Sees

We do not give the agent the latent internal state (*f*_*t*_, *s*_1,*t*_, *s*_2,*t*_). Instead, the agent observes a short window of past actions and errors, **o**_*t*_, where

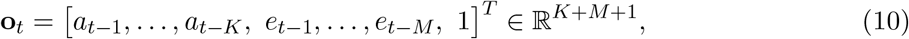

where *K* and *M* are the context- and error-history depths, and the trailing 1 is a bias term. This is a deliberately limited observation. It is chosen to mirror what a teacher could plausibly observe without a formal model of the learner. A clinician sees the patient’s recent attempts and outcomes; a coach sees the recent shots and where they landed. The latent components of memory are not directly accessible in reality, so we do not expose them to the RL agent. We also deliberately omit an explicit time-to-go signal (*T* − *t*)*/T*. The agent recovers the time-structured block–repair– interleave motif (Section 9) from behavioral history alone, and whether an explicit time-to-go input removes the high-*ρ* terminal-switch collapse (Section 11) is left to future work.

The choice of (*K, M*) controls how much behavioral history the agent can use. We treat it as an experimental variable in Section 10, where we ask how much memory is sufficient to learn near-optimal context schedules.

### 7.2 Algorithms

We compare two algorithms, REINFORCE (Williams, 1992) and PPO (Schulman et al., 2017). Our REINFORCE model consists of a linear softmax policy on *o*_*t*_ trained with discounted returns and a baseline. By contrast, our PPO model consists of a linear actor and a small two-layer critic, trained with the standard clipped surrogate objective and Generalized Advantage Estimation. As REINFORCE models typically suffer from high variance, it is used as a simple control, while PPO is the primary RL model of focus.

All agents are trained with 1,000 episodes of *T* = 20 trials by default; long-horizon agents use *T* = 200. We deliberately keep the architectures small. Our goal is to determine what kind of policy is learnable, not how fast or large a network can be made to fit.

### 7.3 Per-Objective Training

For every combination of session length *T* and tradeoff parameter *ρ*, we train a separate reinforcement learning agent. This “specialist” approach is a deliberate choice and our goal is to evaluate whether RL can discover the optimal schedule for a specific, fixed objective, directly matching the exact solver’s solution at that point. We do not attempt to train a single “generalist” agent that can generalize across different acquisition-retention tradeoffs. Designing policies that can dynamically adjust to varying objectives introduces separate methodological challenges (such as multi-task or meta-reinforcement learning) which are outside the scope of this benchmark.

## 8 The Agent Recovers the Exact Optimum

Figure 5 provides the total loss for a *T* = 20 session across a range of *ρ* values for *ρ* ∈ [0.01, 0.07], in which separately trained PPO and REINFORCE models are evaluated at each value of *ρ* against the exact optimum and the static baselines. Given a 5 × 5 (context history length × error history length) history window, PPO matches the exact optimum to within the small-*ρ* alternation gap (1.118 vs. 1.102 at *ρ* = 0.01; see Section 6) and matches it across the remainder of the sweep, including the mixed-optimum interior. Restricted to a 1 × 1 window, PPO still matches in the alternating regime but drifts modestly past the phase boundary, where recovering the optimum requires recognizing the late-corrective-switch opportunity from accumulated state that a single-step window cannot represent. REINFORCE, by contrast, is noticeably less stable in the high-*ρ* regime. The 1 × 1 variant flips between alternating and blocked-with-switch behavior, while the 2 × 2 variant collapses to alternating. We attribute this to the high variance of the unclipped policy gradient, which prevents reliable convergence to the late-switch family.

**Figure 5:**
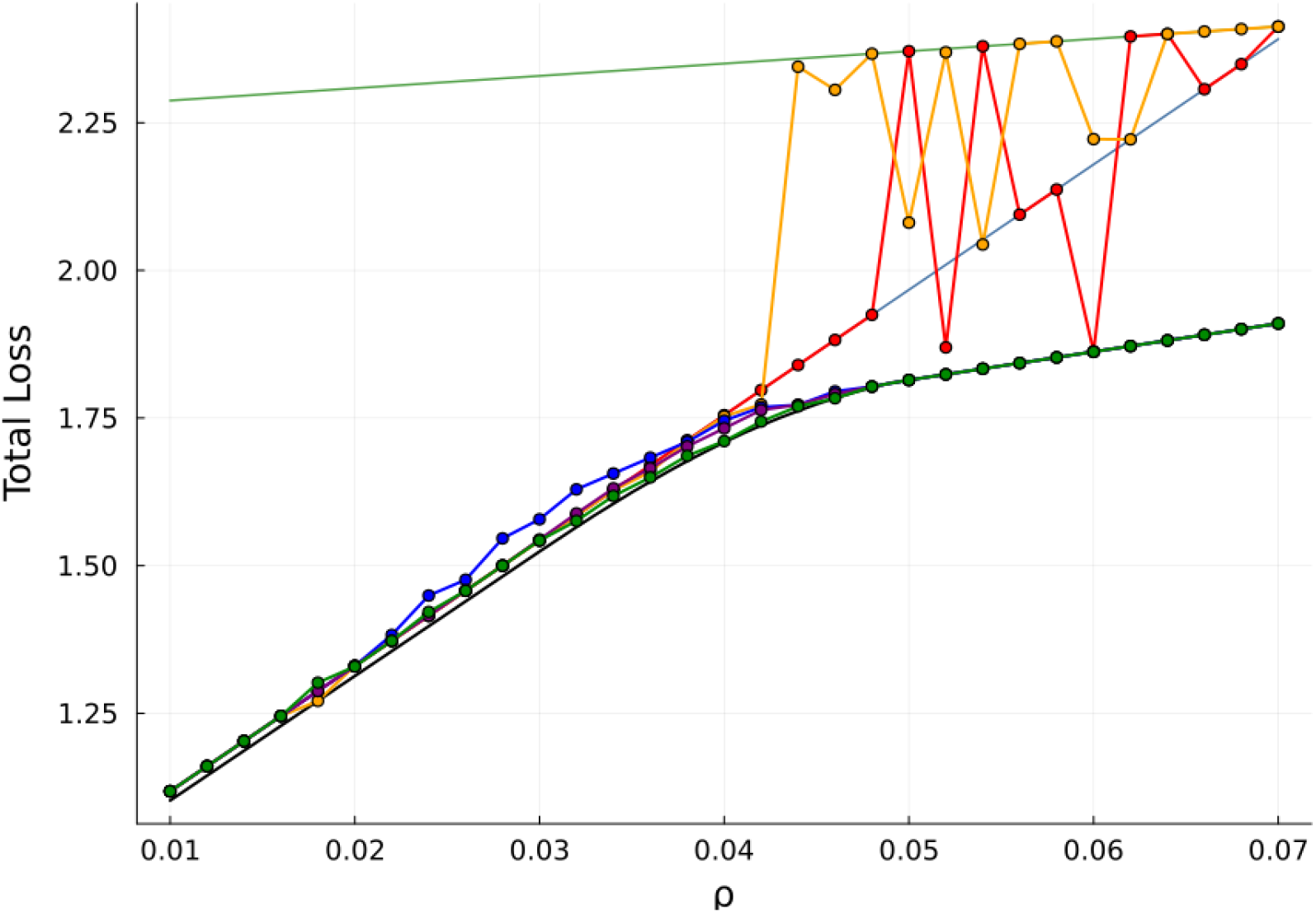
RL scheduler comparison for horizon *T* = 20 trials. The black curve represents the exact optimum; the alternating and blocked baselines are plotted in light blue and light green, respectively. Markers indicate the reinforcement learning policies: PPO (1 × 1 in blue, 2 × 2 in purple, and 5 × 5 in dark green) and REINFORCE (1 × 1 in red and 2 × 2 in orange). Per-*ρ* PPO with sufficient history (5 × 5, dark green) matches the exact optimum across the sweep *ρ* ∈ [0.01, 0.07]: exactly through the mixed-optimum interior, and to within the small-*ρ* terminal-correction gap (1.118 vs. 1.102 at *ρ* = 0.01). Smaller-history PPO (1 × 1 and 2 × 2) matches in the alternating regime but drifts modestly past the phase boundary, where recovering the optimum requires recognizing the late-corrective-switch opportunity from accumulated state. REINFORCE models exhibit high instability in the high-*ρ* regime, which we attribute to the high variance of the unclipped policy gradient preventing convergence to the required late-corrective-switch structure.

None of this is surprising on its own; it is what we would expect from each algorithm on a problem of this type. The interesting fact is that PPO with sufficient history matches the exact optimum across the *ρ* sweep: exactly in the mixed-optimum interior, where the optimal schedule is not an element of the static endpoint family, and to within the 0.016 small-*ρ* terminal-correction gap in the retention-dominated regime. The agent observes only past actions and errors, with no direct knowledge of the latent fast–slow state, and still recovers the right schedule for each regime.

## 9 Long-Horizon Recovery and the Block–Repair–Interleave Motif

Figure 6 reports the *T* = 200 sweep for the specialist agents alongside the static baselines and beam search. The plot includes both the alternating baseline (rising linearly with *ρ*) and the blocked-plus-terminal-switch baseline (dashed gray, rising slowly), the latter being the relevant asymptote for the upper end of the sweep. The two static curves *cross* between *ρ* = 0.0125 and *ρ* = 0.015: alternation dominates below the crossover, blocked-plus-terminal-switch dominates above. The largest gap between the best static baseline and beam — which is the most useful definition of “sweet spot” — is concentrated at the crossover and reaches roughly 0.49 at *ρ* ≈ 0.015.

**Figure 6:**
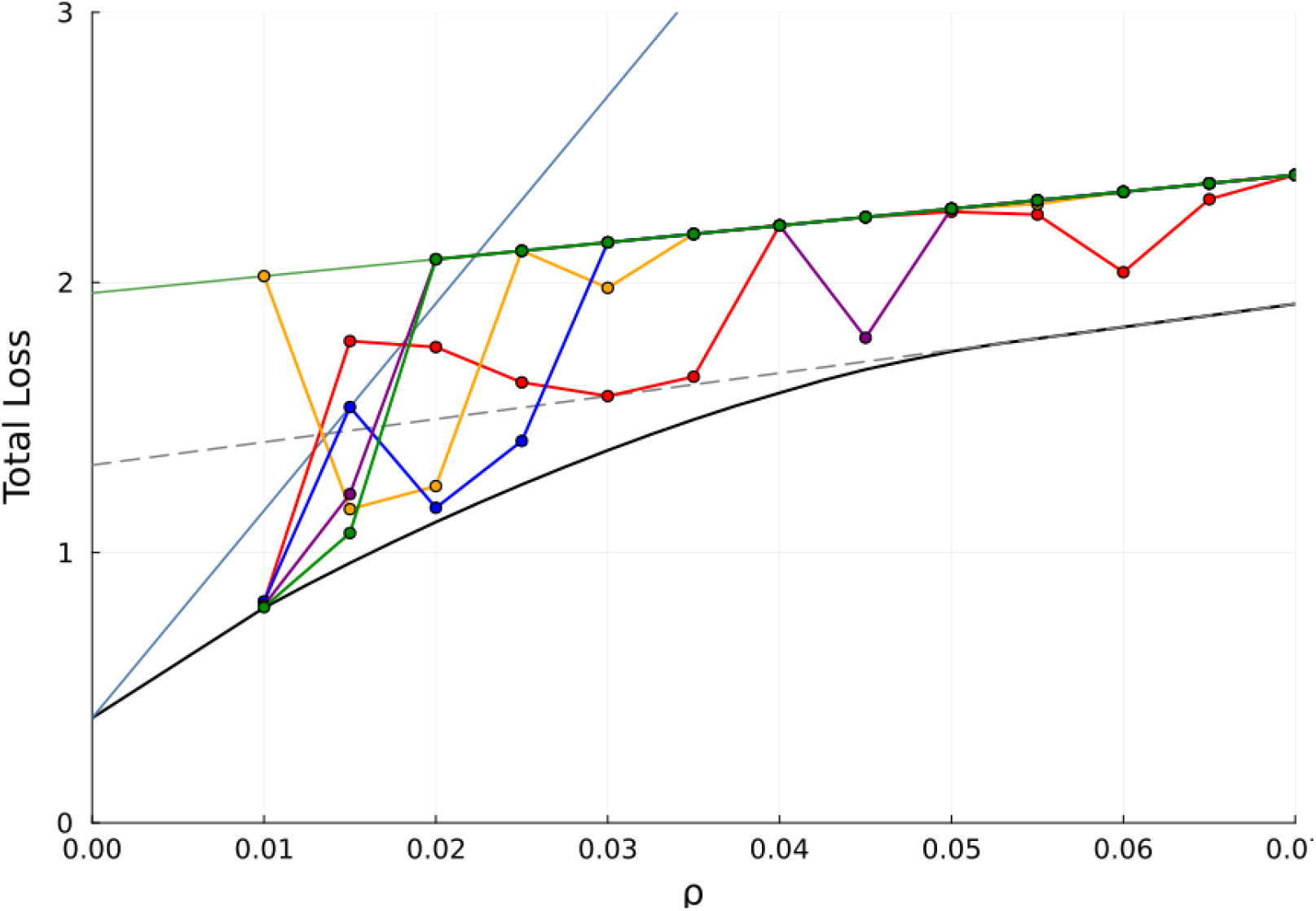
RL scheduler comparison for a long horizon session (*T* = 200 trials). The black curve represents the beam-search upper bound; the alternating and blocked baselines are plotted in light blue and light green, respectively, and the blocked-plus-terminal-switch baseline is plotted in dashed gray. Markers indicate the reinforcement learning policies: PPO (1 × 1 in blue, 2 × 2 in purple, and 5 × 5 in dark green) and REINFORCE (1 × 1 in red and 2 × 2 in orange). The two relevant static asymptotes cross near *ρ* ≈ 0.014, and the largest gap between the best static baseline and beam is concentrated near that crossover. PPO specialists with sufficient history (5 × 5, dark green) close most of the gap in *ρ* ∈ [0.01, 0.018]; above that, blocked-plus-switch is itself close to beam, but the PPO specialists collapse to *pure* blocked rather than blocked-plus-switch and miss the easy late-switch fix.

In this sweet-spot interval, PPO with sufficient history closes most of that gap. At *ρ* = 0.01, PPO 5 × 5 achieves total loss 0.798, beam achieves 0.796, and the better of the two relevant static baselines (alternation here) is 1.155. At *ρ* = 0.015, PPO 5 × 5 achieves 1.072, beam achieves 0.962, and the better static baseline (blocked-plus-switch) is 1.452. Above *ρ* ≈ 0.018 the picture changes qualitatively. Blocked-plus-terminal-switch is itself within 0.07 of beam at *ρ* = 0.04 (1.665 vs. 1.591), so the headroom for an RL policy is small. The PPO specialists in this regime, however, do not even reach blocked-plus-switch; they collapse to pure blocking and miss the easy late-switch fix entirely. We treat this as a genuine failure mode in Section 11.

The interesting result is not the loss curves, but the schedules behind them. Figure 7 shows learned schedules at the boundaries of the regime. At *T* = 200, *ρ* = 0.01, the agent reproduces the block–repair–interleave macrostructure that beam search converges to (shown in Fig. 4). The visual identity of the two structures is, in our view, the strongest single piece of evidence in the paper that the agent is learning the right kind of policy — not just a loss-tracking curve, but a schedule with the same teacher’s instinct.

**Figure 7:**
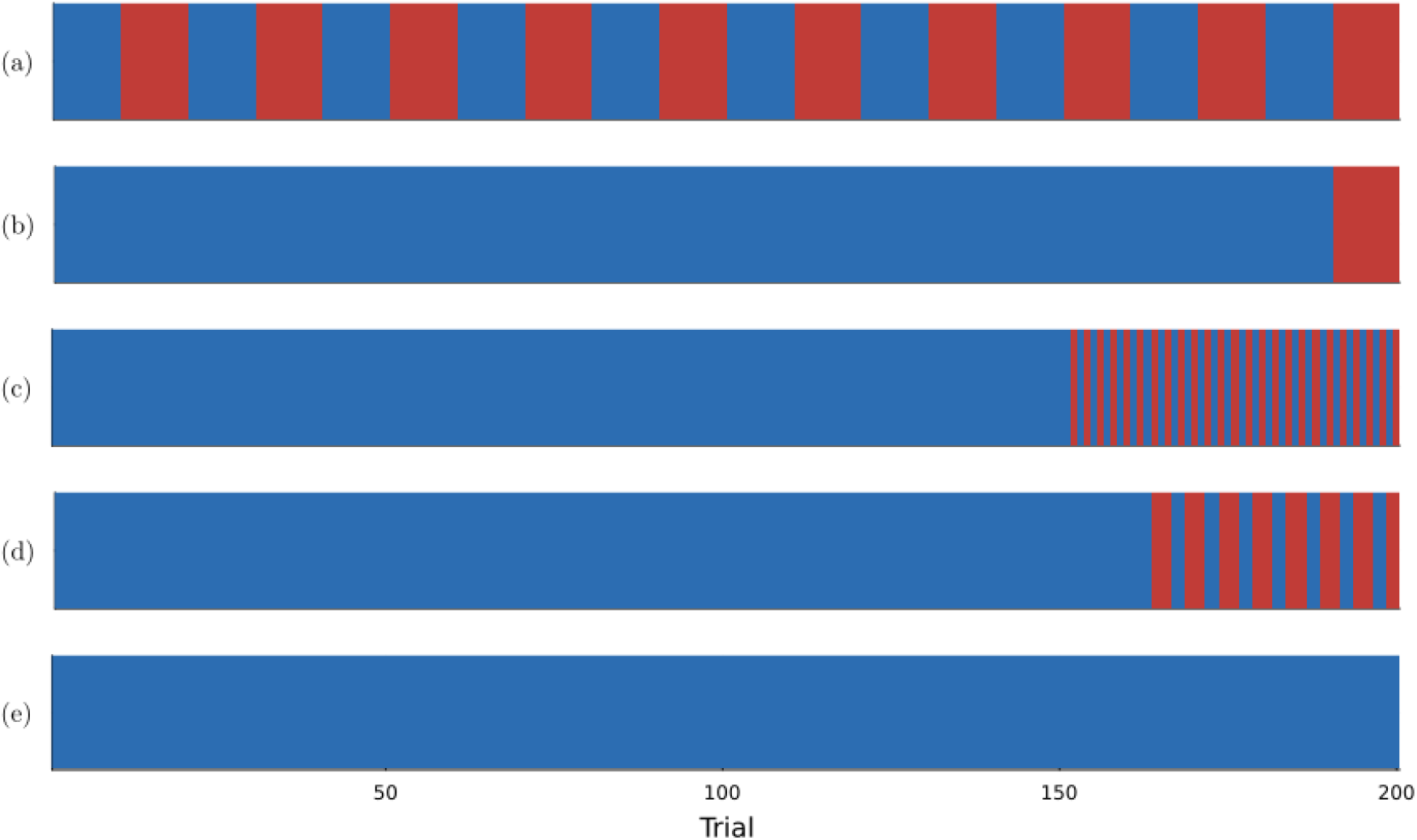
Learned PPO (5 × 5) schedules across the regimes. (a) *T* = 20, *ρ* = 0.01, loss = 1.118: strict alternation. (b) *T* = 20, *ρ* = 0.05, loss = 1.814: blocked practice plus a terminal switch. (c) *T* = 200, *ρ* = 0.01, loss = 0.798: the block–repair–interleave motif (long exploitation block, transition, interleaved tail; compare Figure 4). (d) *T* = 200, *ρ* = 0.015, loss = 1.072: a cleaner BRI structure near the sweet spot, with a longer block and a more compact interleaved tail. (e) *T* = 200, *ρ* = 0.04, loss = 2.211: the policy has collapsed to pure blocking with no terminal switch (where blocked-plus-terminal-switch achieves 1.665 and beam achieves 1.591, leaving a single-bit edit on the table; see failure mode discussed in Section 11).

The block–repair–interleave motif is sharpest at *ρ* = 0.015 (panel d): an initial block of about 165 trials in one context, a brief transition, and a compact interleaved tail of about 35 trials near the probe. The same agent collapses to pure blocking somewhere between *ρ* = 0.015 and *ρ* = 0.02 and stays there for the rest of the sweep (panel e shows the representative *ρ* = 0.04, where blocked-plus-terminal-switch is itself within 0.07 of beam). We discuss this transition further as a failure mode in Section 11.

## 10 What Memory Does the Teacher Need?

Because the teacher operates under partial observability, its policy relies entirely on the learner’s behavioral history of past actions and errors. How much history is necessary to recover the optimal schedule? This question is central to the framework’s practical applicability. In real-world settings, a teacher cannot access a learner’s latent memory states and must direct practice using only these behavioral signals.

We systematically sweep the context- and error-history depths *K, M* ∈ {1, …, 20} on a 20 × 20 grid, training a separate PPO agent at each cell across three random seeds with a training budget scaled to accommodate the larger observation dimensions. Figure 8 shows the resulting total loss and switch rate across this sweep for *T* = 20 and *ρ* = 0.01.

**Figure 8:**
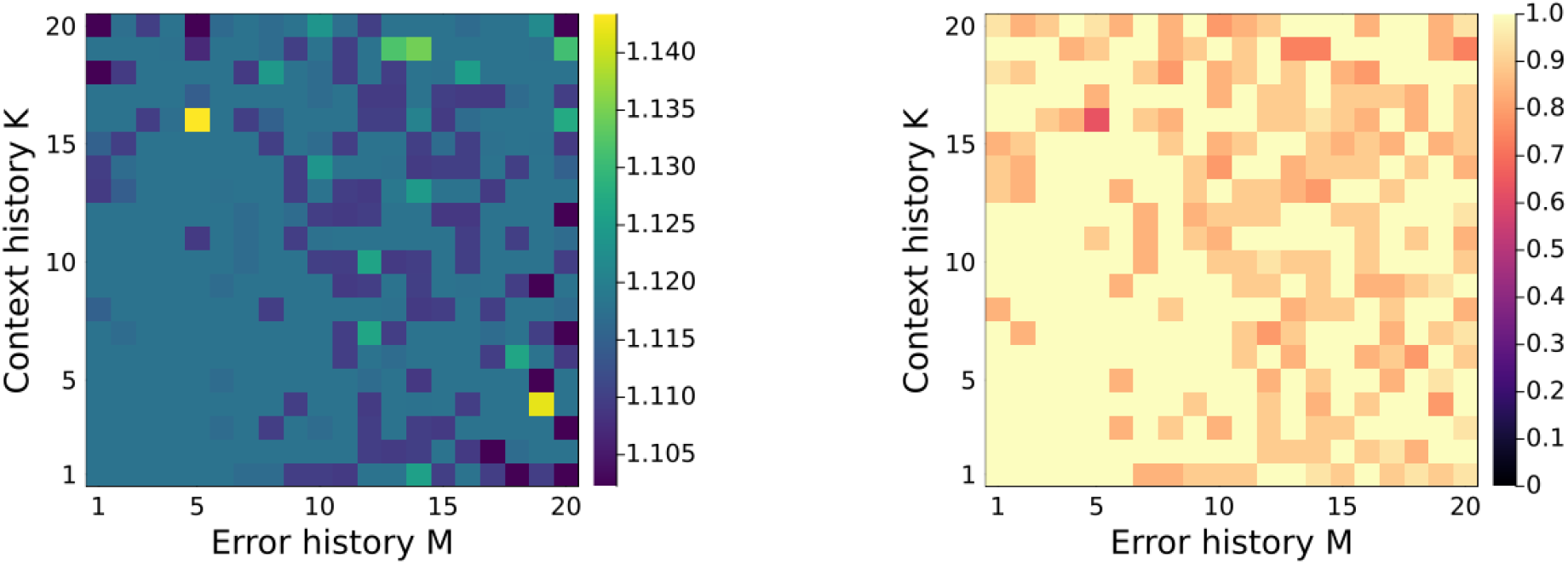
PPO history-depth hyperparameter sweep at *T* = 20, *ρ* = 0.01, showing total loss (left panel) and switch rate (right panel), where colorbar intensity represents total loss and switch rate respectively. The teacher does not need to see the latent state; every history configuration on the 20 × 20 grid recovers near-exact loss (range 1.10–1.14 across the grid; exact 1.102). The right panel (switch rate) shows that most configurations learn strict alternation (switch rate 1.0), with a grid-mean of 0.947 that coincides with the exact optimum’s switch rate (0.947); the agents thus sit at or just above the optimum, missing only its single terminal correction. The handful of bright cells in the loss panel are isolated bad seeds rather than a structured failure region.

These results demonstrate that the teacher does not require access to the learner’s latent memory states. A small behavioral window of past actions and errors is sufficient to recover near-optimal schedules, indicating that the latent fast–slow dynamics are reconstructible from behavior at these time scales.

### 10.1 Sufficiency of History in the Mixed Regime

While the initial hyperparameter sweep (Figure 8) focuses on the alternation-dominated regime (*ρ* = 0.01), this is the simplest structure to discover. A more challenging question is whether a similarly modest history window is sufficient in the mixed-regime interior, where optimal schedules exhibit significant state-dependent structure. To test this, we repeat the 20 × 20 sweep at *ρ* = 0.04 (the mixed-regime interior for *T* = 20, with an exact loss of 1.709 and an exact switch rate of 0.474). The results are presented in Figure 9.

**Figure 9:**
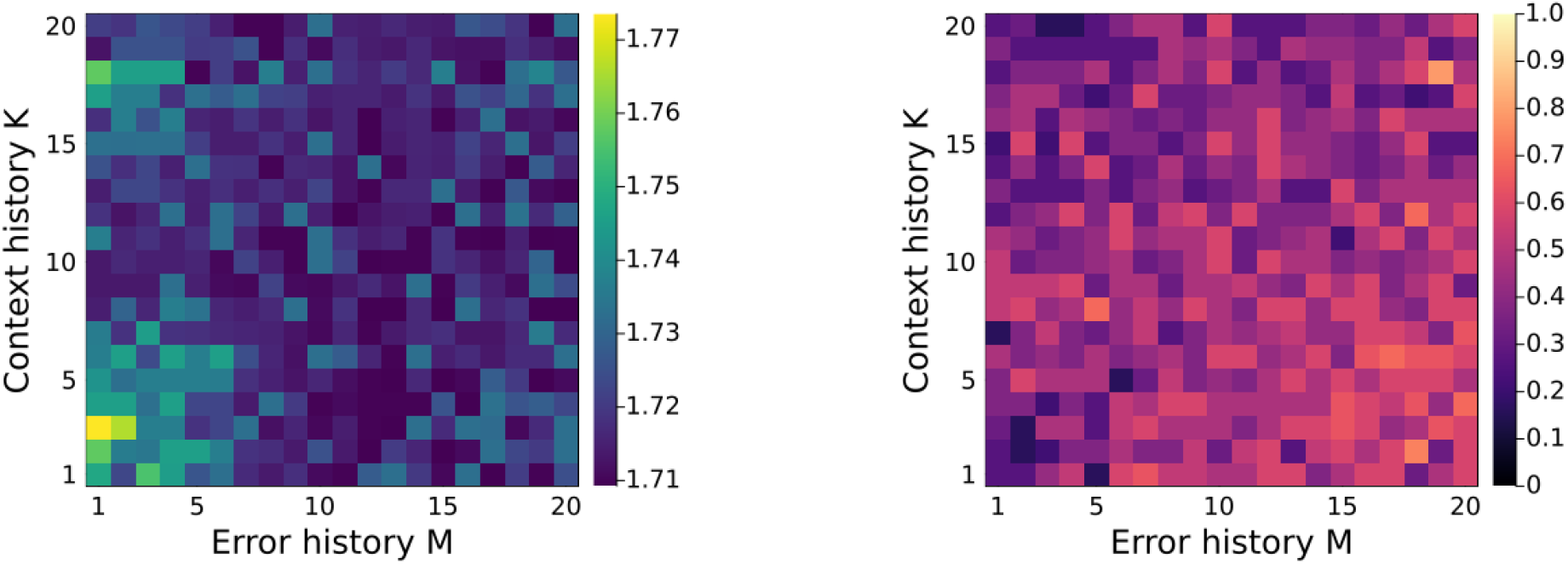
PPO history-depth hyperparameter sweep at *T* = 20, *ρ* = 0.04 (mixed regime), showing total loss (left panel) and switch rate (right panel), where colorbar intensity represents total loss and switch rate respectively. Loss ranges 1.71–1.77 across the grid against exact 1.709; the worst cells are at low error-history depth *M* where the input is too thin to disambiguate the mid-regime state. The switch-rate panel hovers near the exact value 0.474 across most of the grid. The primary result of Figure 8, that modest behavioral history is sufficient, holds in the harder mixed regime as well, with no qualitative change.

Taken together, these two hyperparameter sweeps support our broader claim. Whether the optimal policy is at or near strict alternation (*ρ* = 0.01) or a complex, state-dependent mixed schedule (*ρ* = 0.04), a short behavioral history window of past actions and errors enables PPO to achieve near-optimal performance. At this short horizon, the agent recovers the optimal schedules without requiring privileged access to the learner’s latent fast–slow states in either regime.

## 11 What the Agent Cannot Do

The exact solver and the beam heuristic allow us to analyze precisely where and why the reinforcement learning agents fall short. We document three distinct failure modes: (1) a sharp collapse to pure blocked practice in the high-*ρ* (performance-dominated) regime, failing to recover a simple terminal-switch correction; (2) severe optimization instability in REINFORCE near regime boundaries due to high policy-gradient variance; and (3) a small residual gap in the intermediate sweet-spot regime where PPO recovers the qualitative block–repair–interleave macrostructure but transitions between phases suboptimally.

### 11.1 High-*ρ* Collapse and the Missing Terminal Switch

For long-horizon sessions (*T* = 200), the theoretical ceiling above the sweet spot is set by the blocked-plus-terminal-switch baseline (which sits within 0.07 of the beam-search upper bound at *ρ* = 0.04). Remarkably, PPO fails to reach even this baseline. While the 5 × 5 PPO agent recovers a clean block–repair–interleave structure at *ρ* = 0.015 (Figure 7d, achieving a loss of 1.072 against the beam-search upper bound of 0.962), increasing the tradeoff parameter slightly to *ρ* = 0.020 causes the policy to collapse to pure blocking (yielding a loss of 2.086). By *ρ* = 0.040, the agent’s schedule becomes a solid, uncorrected blocked block (Figure 7e) with a total loss of 2.211—identical to the pure-blocked baseline. In contrast, the blocked-plus-switch baseline achieves 1.665 and the beam-search upper bound achieves 1.591.

This means the agent leaves a straightforward performance improvement on the table: inserting a single, well-placed terminal switch would close most of the gap to the optimum. However, under the standard PPO surrogate objective, the gradient signal for introducing one beneficial switch near the end of a 200-trial session is extremely weak. This transition from the block–repair–interleave motif at *ρ* = 0.015 to a collapsed, purely blocked policy at *ρ* = 0.020 is remarkably sharp. Although smaller-history PPO variants (such as 1 × 1 and 2 × 2) occasionally discover better schedules at intermediate values like *ρ* = 0.020, the same collapse to pure blocking eventually re-emerges as *ρ* increases.

#### 11.1.1 Diagnostic Probe: Warm-Starting from Blocked-Plus-Switch

To explain this high-*ρ* failure, we consider two hypotheses: (1) an exploration failure, where PPO initialized from scratch cannot discover a single beneficial switch near the end of a 200-trial session; and (2) an optimization failure, where the surrogate gradient surface near the blocked-plus-switch schedule does not support it as a stable attractor. To identify the failure mode, we design a warm-start diagnostic probe at (*T, ρ*) = (200, 0.04). We create the target blocked-plus-terminal-switch schedule and behavior-clone a PPO actor to mimic it (using cross-entropy loss over 32 noisy trajectories for 2, 000 epochs). After confirming that the warm-started actor successfully reproduces the blocked-plus-switch policy with a deterministic rollout loss of 1.665, we resume standard PPO training for an additional 5, 000 episodes. Figure 10 tracks the deterministic rollout loss every 25 policy iterations.

**Figure 10:**
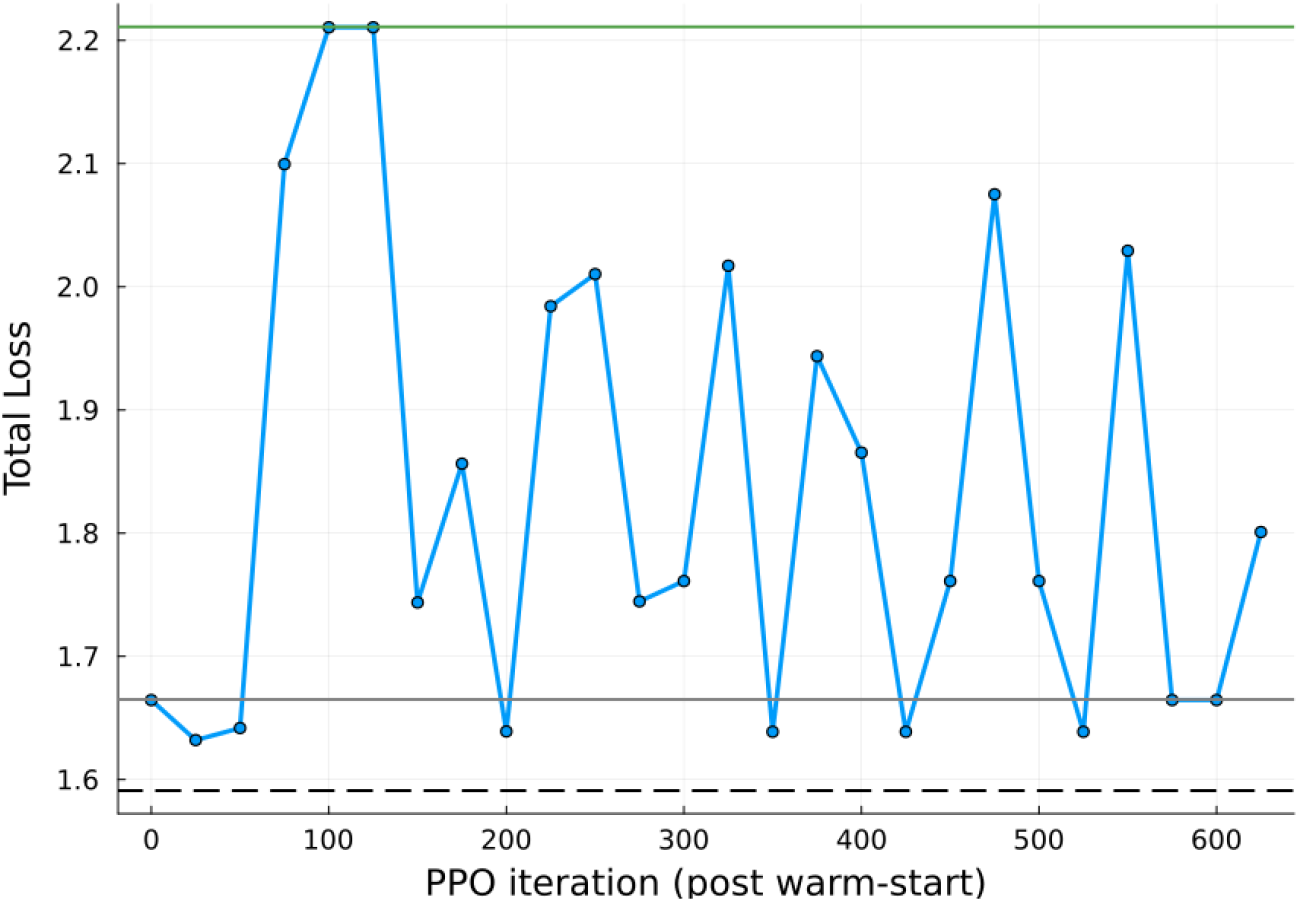
Warm-starting PPO from blocked-plus-terminal-switch at *T* = 200, *ρ* = 0.04, showing the trajectory of the policy’s rollout loss (solid blue curve with markers) during continuation training. Reference baselines are shown as horizontal lines: beam search at 1.591 (dashed black), blocked-plus-last-switch at 1.665 (solid gray), and pure blocked at 2.211 (solid green). The deterministic rollout begins at the warm-start loss of 1.665 (matching blocked-plus-switch) and oscillates between two attractors throughout the continuation: blocked-plus-switch (loss ≈ 1.65, terminal action 2) and pure blocked (loss 2.211, terminal action 1). Neither basin is stable under PPO’s gradient at this *ρ*; the policy never settles into the better of the two.

The resulting trajectory rules out a pure exploration failure, as the policy successfully retains the target behavior for several iterations. It also rules out a simple optimization failure, since the policy does not immediately degenerate to pure blocking. Instead, the system exhibits metastability. Both blocked-plus-switch and pure blocking act as local attractors in the surrogate gradient field at *ρ* = 0.04, and policy-gradient stochasticity drives the deterministic rollout back and forth between them. Because the presence of a terminal switch is essentially a single-bit edit in the schedule, the transition between these two states is discrete. Whether the agent loses the switch (drifting to pure blocking) or recovers it (returning to blocked-plus-switch) depends entirely on whether the policy’s argmax action flips on a single terminal trial. The PPO objective does not provide a sufficiently strong gradient to anchor the policy at the better of the two attractors, resulting in indefinite oscillation. This highlights a fundamental limitation of standard policy-gradient methods when applied to schedule optimization. When crucial corrections depend on sparse, binary decisions, the optimization landscape becomes near-degenerate, and standard surrogate objectives lack the mechanism to break the symmetry and stabilize the optimal sparse switch.

### 11.2 REINFORCE Variance

REINFORCE achieves comparable losses to PPO in the easy regimes (alternation and pure blocking are both stable attractors) but is conspicuously worse in the mixed and blocked-with-switch regimes (Figure 5). REINFORCE 1 × 1 exhibits stochastic alternating–blocked flips above *ρ* = 0.04, and REINFORCE 2 × 2 collapses to alternating well past the phase boundary. We attribute the gap to PPO’s clipped surrogate stabilizing high-variance gradients near the boundaries between schedule families.

### 11.3 Residual Sweet-Spot Gap

At *ρ* = 0.015 (the middle of the long-horizon sweet spot), the best PPO specialist achieves 1.072 while beam search achieves 0.962, a gap of about 11% of the loss. This regime requires the schedule to execute a finely tuned block–repair–interleave with the transition placed at a specific point. Gradient methods recover the macrostructure but place the transition slightly suboptimally. We have not closed this gap.

We return in the discussion (Section 12) to the broader point that the exact solver and the beam heuristic are what make these failures legible at all.

## 12 Discussion

We have framed practice scheduling as a sequential decision-making problem on a dual-rate motor learner, characterized the optimal-schedule family in a minimal benchmark, and shown that a model-free reinforcement-learning agent can recover this family purely through trial-and-error interaction. We close by tracing the framework’s implications back to the literatures it touches.

### 12.1 Contextual Interference, Reconsidered

The contextual interference (CI) effect — blocked practice favoring acquisition, interleaved practice favoring retention — is a robust empirical finding (Schmidt and Bjork, 1992; Guadagnoli and Lee, 2004; Czyż et al., 2024) that has been interpreted as a property of practice schedules. In our framework, the dissociation it describes is not a property of schedules but a consequence of the learner’s memory architecture. A fast process favors local exploitation; slow context-specific processes favor distributed practice. When acquisition is the only objective, the fast-process bias dominates and blocked-with-correction practice wins. When retention is the only objective, the slow-process bias dominates and (near-)alternating practice wins. In between, which is where most real practice lives, the optimum is neither, and the right schedule is a mixed one whose structure depends on the learner’s current state.

This reframing does not contradict the empirical CI literature; it explains it. The endpoint regimes of the learner (alternating when the acquisition-retention tradeoff parameter *ρ* is small, blocked-with-correction when it is large) are exactly the regimes in which a CI-style experiment would be set up. The mixed regime is the regime in which the experimental literature, designed around fixed schedule families, has historically been silent. The phase diagram of Figure 3 suggests that this regime is where the most interesting empirical work remains to be done.

### 12.2 Implications for Motor Learning Experiments

We do not claim that the learner is the right model for any particular motor adaptation task. We do claim that the framework (exact-optima as ground truth, schedule families as a *ρ*-indexed manifold, learned policy as a recovery test) is a useful template for studies that use richer learners. Three concrete predictions arise from the learner and are worth verifying against experimental data on dual-rate adaptation models that have been fit to behavior:

1. Within a given task family and learner population, the optimal schedule should depend on the experimentally specified weighting of within-session performance against retention. Studies that vary this weighting (for example by adjusting the delay Δ to the probe, or by changing the relative reward of within-session and probe accuracy) should observe optimal schedules transitioning through the three regimes we describe.
2. In long sessions, a block–repair–interleave structure should outperform both blocked and interleaved practice in the intermediate weighting regime. Schedules consisting of a single early block, a brief switch, and an interleaved tail are not standard experimental conditions; they are predicted to be the optimal ones in a defined window.
3. A behavioral teacher, human or learned, should be able to recover schedules of approximately optimal structure without access to the learner’s internal state. Modest histories of past trials and outcomes should suffice. This is our methodological prediction, and it is the place where the RL contribution makes contact with how a real teacher operates.

### 12.3 Implications for Finite-Time Practice Scheduling

The most direct practical analogy is finite-time practice within a single session. A clinician working with a patient for 30 to 45 minutes is in approximately the long-horizon regime relative to the learner’s time scales, and the patient’s retention is the dominant outcome of interest. This describes a small-*ρ* regime. The learner’s prediction in that regime is unambiguous, the schedule should not be blocked, but neither should it be strict alternation. The optimal structure has an early exploitation block (which the practitioner often calls a warm-up), a corrective transition (the moment when the practitioner switches to a neglected target), and a sustained interleaved tail near the session’s end.

We are aware that this is a coarse mapping. The learner is not a model of motor recovery, the patient population in any rehabilitation setting is heterogeneous, and the actual time scales of fast and slow adaptation in clinical contexts differ across individuals and tasks. We are not proposing it as a treatment-planning tool. We are proposing it as a way of giving a precise meaning to the practical intuitions that practitioners already use, and as a benchmark against which more behaviorally grounded models can be tested. The methodology of computing the optimal schedule where you can, characterizing its dependence on the objective parameter, and testing whether a learned policy recovers it from realistic observability transfers regardless of whether its specific predictions do.

### 12.4 What the Benchmark Does Not Establish

We are explicit about three limits.

- **One learner type**. All results are for the fast–slow learner with two contexts. We confirm in Appendix B that the three-regime phase structure is preserved across four parameter sets that span the learner’s constrained parameter region; only the locations of the regime boundaries shift. We do not establish that the phase structure survives in richer architectures (more contexts, stochastic dynamics, hierarchical memory). A *K* = 3-context extension is the natural next experiment but is out of scope here: the brute-force backend grows as *K*^*T*^ and would require Mixed-Integer Programming (MIP) formulations (Wolsey, 1998) to remain tractable, while “alternation” itself becomes multi-valued.
- **Open-loop objective**. The reward and the evaluation use the same open-loop loss (1). A teacher uncertain of the model (i.e., not knowing which of several learner-parameter regimes the patient occupies) faces a more difficult problem. Bayesian or risk-aware extensions are a natural next step.
- **Synthetic learner**. The learner is not fit to behavioral data. Its parameters were chosen to make the manifold structure visible, not to match a particular task. The framework’s value is methodological. Whether the framework should be ported to a behaviorally fit model is an empirical question for the dual-rate motor-learning community.

### 12.5 Why Benchmarks Like This Matter for Empirical RL

A separate methodological observation closes the discussion. The specific failure modes we report — the high-*ρ* ceiling collapse, the residual sweet-spot gap, the REINFORCE variance — are visible only because the exact solver and the beam heuristic give us a non-RL ground truth to compare against. Without that ground truth, the same numbers would read as “RL works approximately well.” We take this as an argument for the value of small benchmarks with computable optima as scaffolding for empirical-RL methodology, independent of this specific learner.

## 13 Conclusion

We posed the question “when should a teacher block, interleave, or do something else?” in a minimal fast–slow learner whose optimal practice schedules can be computed exactly. The answer is a *ρ*-indexed family of schedules with three regimes and a long-session sweet spot in which the optimum has an interpretable block–repair–interleave structure. A reinforcement-learning agent observing only the learner’s behavior recovers this family from interaction. The exact solver lets us audit the comparison: where the agent matches the optimum, where it falls short, and what kind of memory it needs.

The contextual interference effect, which has driven decades of motor-learning research, is the boundary case of a richer state-dependent control problem. The block–repair–interleave structure that emerges in the sweet spot is the prediction the learner makes about how a finite practice session should be organized when retention is the dominant outcome of interest. Whether the prediction holds in richer learners and in real practice settings is the empirical question this framework is meant to make precise.

## Code and Data Availability

All code supporting this work — the brute-force exact backend, the beam search, and the full reinforcement-learning training and evaluation pipeline — is publicly archived (Jeter et al., 2026) (active development version at https://github.com/russell-jeter/optimal-practice-schedules-in-a-dual-rate-model-of-motor-adaptation), together with the pretrained PPO and REINFORCE models used to generate the figures, the randomization seeds, training-budget settings, evaluation protocols, and the data underlying every figure. The archive also includes a mixed-integer reformulation with Gurobi and SCIP backends for sessions beyond *T* = 22; it was not used for any result reported here.

## Acknowledgments

This work was supported by a Brains & Behavior Seed Grant from Georgia State University.

## A Beam-search hyperparameters and search space sensitivity analysis

The default beam configuration uses width 16, elite width 4, sample temperature 0.05, and 64 random restarts. Candidate suffixes drawn from each restart: alternation, pure blocking, single-switch schedules, and greedy last-trial rollouts. Diversity selection across the top 20 candidates uses a normalized-Hamming threshold of 0.15. Local refinement applies single-flip and switch-boundary moves until no improvement is found. The solver implementation and the exact configuration used to produce the figures are available in the archived code release (Jeter et al., 2026).

To establish that the block–repair–interleave structure is induced by the objective rather than by a particular search heuristic, we test the sensitivity of the beam search configuration and confirm the qualitative macrostructure survives every modification we tested (Figure 11).

**Figure 11:**
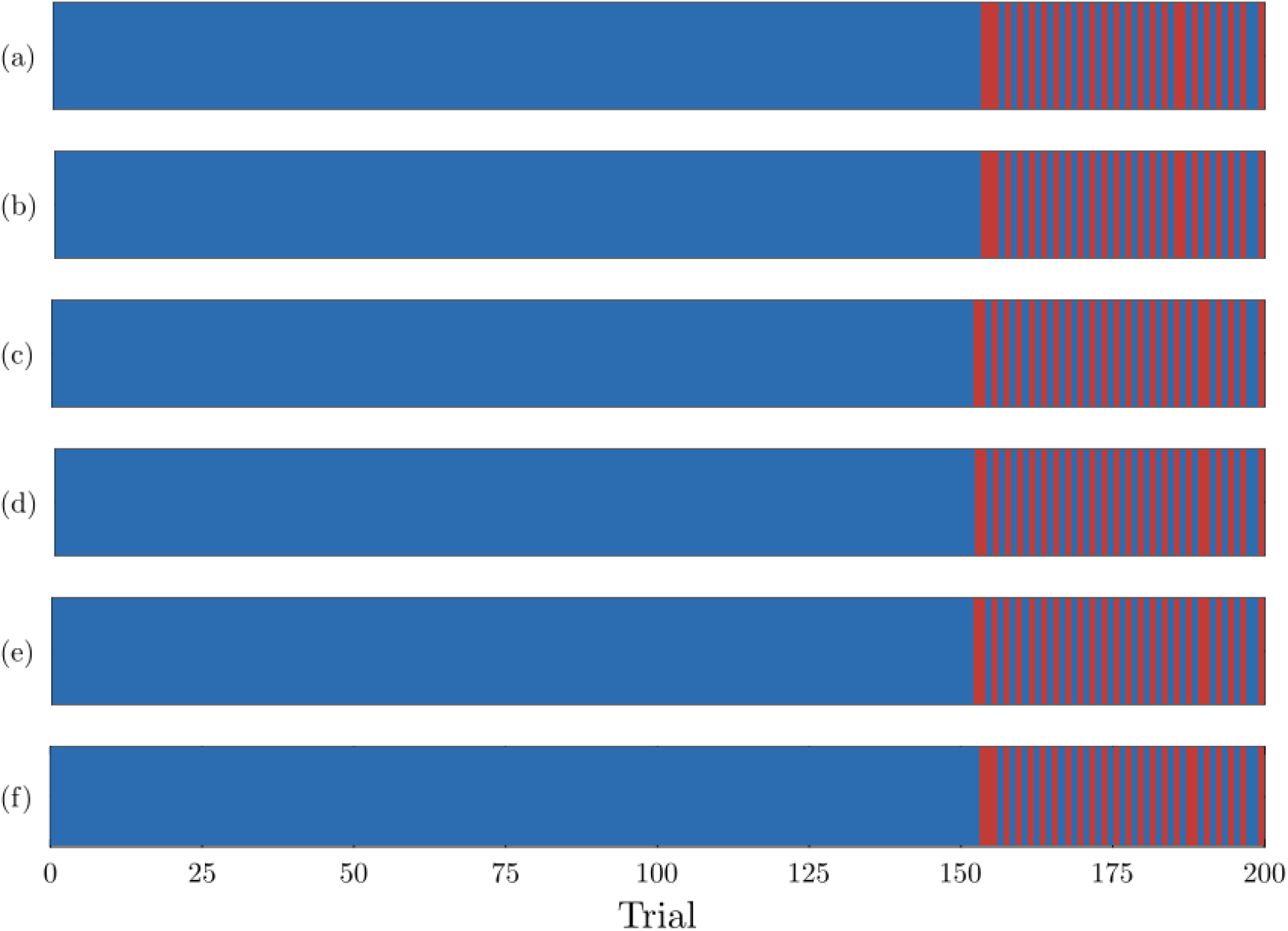
Beam-search sensitivity analysis at (*T, ρ*) = (200, 0.013) across six variants: (a) Full sampled beam (loss = 0.8964, *d* = 0.00), (b) No fallback suffix family (loss = 0.8964, *d* = 0.00), (c) No Prop1 suffix family (loss = 0.8969, *d* = 0.03), (d) No single-switch suffix family (loss = 0.8969, *d* = 0.03), (e) Core heuristics only (loss = 0.8969, *d* = 0.03), and (f) Novelty-pruned beam search (loss = 0.8967, *d* = 0.01), where *d* is the normalized Hamming distance to the full beam-search schedule. Removing any single candidate-suffix family from the search, restricting the search to core heuristics only, or applying novelty pressure to diversify the candidate pool changes details of the recovered schedule but preserves the qualitative block–repair–interleave macrostructure.

## B Parameter robustness

To test whether the three-regime phase structure of Section 6 depends on the gap-amplifying parameter defaults used in the rest of the paper, we regenerate the horizon–*ρ* phase diagram (Figure 3) at four parameter sets that all satisfy the learner’s constraints 0 *< a*_*f*_ *< a*_*s*_ *<* 1 and *b*_*f*_ *> b*_*s*_ *>* 0 but span the constrained region differently:

| Set | $a_f$ | $b_f$ | $a_s$ | $b_s$ |
| --- | --- | --- | --- | --- |
| default | 0.890 | 0.390 | 0.982 | 0.042 |
| less-gap | 0.850 | 0.300 | 0.950 | 0.050 |
| stickier-slow | 0.890 | 0.390 | 0.995 | 0.025 |
| looser-slow | 0.890 | 0.390 | 0.950 | 0.060 |

Figure 12 shows the resulting phase diagrams. The three-regime structure is preserved at every parameter set we tested. The *ρ* values at which the regime transitions occur shift, but in directions that match the dynamics: when slow retention is increased (stickier-slow) the blocked regime expands toward smaller *ρ* because the durable component is harder to lose; when slow retention is decreased and slow learning is faster (looser-slow) the alternating regime expands toward larger *ρ* because the slow process can be repaired cheaply between switches. The qualitative claim of the paper — that the optimal schedule traces a *ρ*-indexed manifold with three regimes, alternating, mixed, and blocked-with-correction — is therefore not an artifact of the paper’s specific defaults.

**Figure 12:**
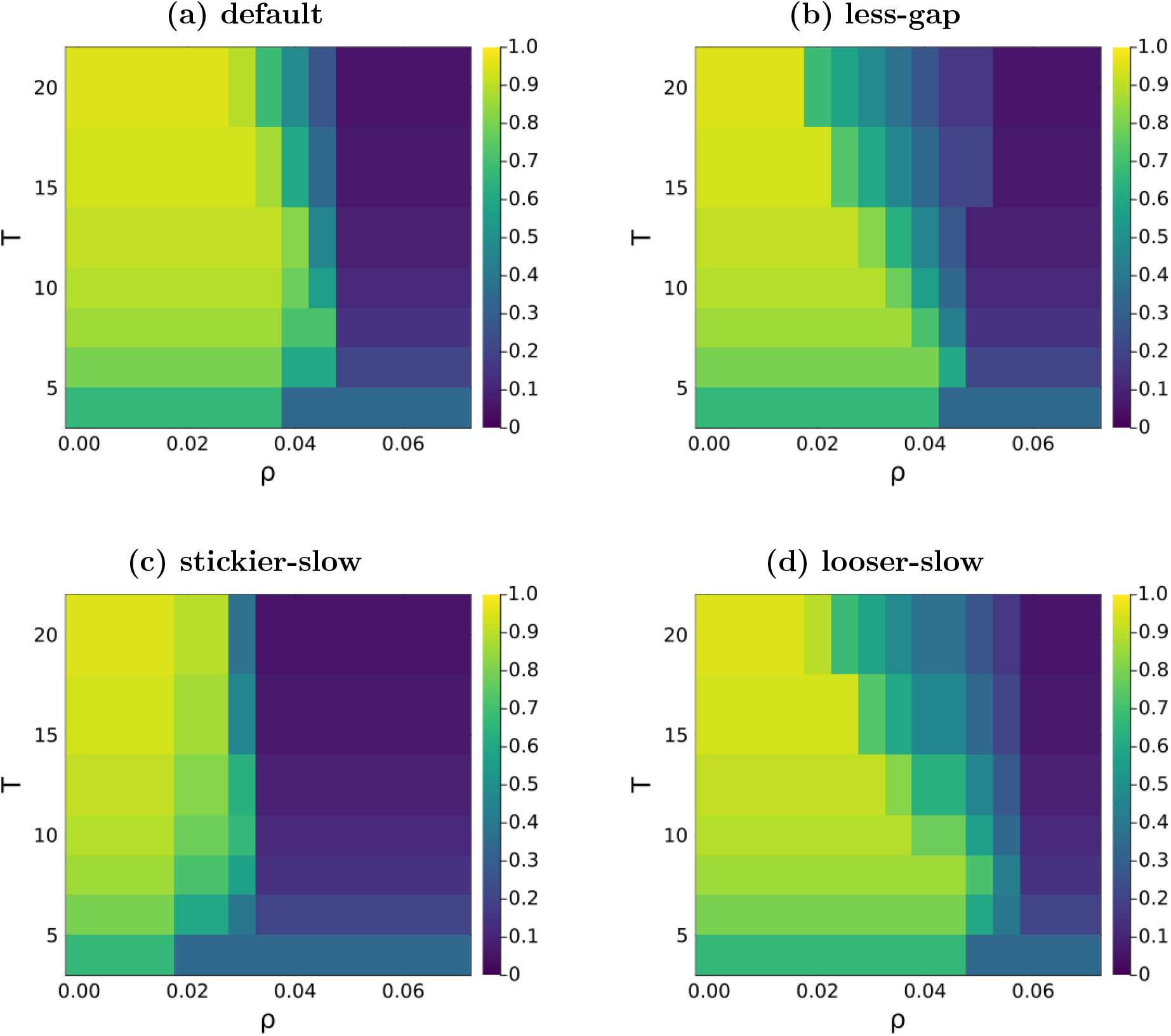
Horizon–*ρ* phase diagrams for four parameter sets: (a) default (*a*_*f*_ = 0.890, *b*_*f*_ = 0.390, *a*_*s*_ = 0.982, *b*_*s*_ = 0.042), (b) less-gap (*a*_*f*_ = 0.850, *b*_*f*_ = 0.300, *a*_*s*_ = 0.950, *b*_*s*_ = 0.050), (c) stickier-slow (*a*_*f*_ = 0.890, *b*_*f*_ = 0.390, *a*_*s*_ = 0.995, *b*_*s*_ = 0.025), and (d) looser-slow (*a*_*f*_ = 0.890, *b*_*f*_ = 0.390, *a*_*s*_ = 0.950, *b*_*s*_ = 0.060). Colorbar intensity represents the switch rate of the exact-optimum schedule. The three-regime structure — high switch rate at small *ρ*, sharp transition at intermediate *ρ*, low switch rate above — is preserved across every set. The location of the transition varies in interpretable ways: stickier-slow has the leftmost crossover (around *ρ* ≈ 0.025, since durable retention is so high that even a small training-error penalty justifies blocking), and looser-slow has the rightmost (around *ρ* ≈ 0.05, since the slow process forgets more between practice sessions and rewards interleaved practice further into the high-*ρ* regime).

## Footnotes

1 Throughout, “blocked” denotes pure single-context practice, meaning all trials in one context, rather than the balanced two-task design with one contiguous block per context that is common in the contextual-interference literature. “Blocked-plus-terminal-switch” is that single-context block with one late repair trial.

## References

Jonathan Bassen, Nikhil Balaji, Michael Schaarschmidt, Candace Thille, Jason Painter, Dawn Zimmaro, and Ethan Fast. Reinforcement learning for the adaptive scheduling of educational activities. In Proceedings of the 2020 CHI Conference on Human Factors in Computing Systems, pages 1–12, 2020. doi: 10.1145/3313831.3376518.

Yoshua Bengio, Jérôme Louradour, Ronan Collobert, and Jason Weston. Curriculum learning. In Proceedings of the 26th International Conference on Machine Learning, pages 41–48, 2009. doi: 10.1145/1553374.1553380.

Max Berniker and Konrad P. Kording. Estimating the relevance of world disturbances to explain savings, interference and long-term motor adaptation effects. PLoS Computational Biology, 7 (10):e1002210, 2011. doi: 10.1371/journal.pcbi.1002210.

Younggeun Choi, Feng Qi, James Gordon, and Nicolas Schweighofer. Performance-based adaptive schedules enhance motor learning. Journal of Motor Behavior, 40(4):273–280, 2008. doi: 10.3200/JMBR.40.4.273-280.

Sebastian H. Czyż, Agata M. Wójcik, Petra Solarská, and Przemyslaw Kiper. High contextual interference improves retention in motor learning: Systematic review and meta-analysis. Scientific Reports, 14:13279, 2024. doi: 10.1038/s41598-024-64206-7.

Mark A. Guadagnoli and Timothy D. Lee. Challenge point: A framework for conceptualizing the effects of various practice conditions in motor learning. Journal of Motor Behavior, 36(2): 212–224, 2004. doi: 10.3200/JMBR.36.2.212-224.

Alkis M Hadjiosif, John W Krakauer, Adrian M Haith, and Maurice A Smith. A double dissociation between savings and long-term memory in motor learning. PLoS Biology, 21(1):e3001928, 2023.

Masahiko Haruno, Daniel M. Wolpert, and Mitsuo Kawato. Mosaic model for sensorimotor learning and control. Neural Computation, 13(10):2201–2220, 2001. doi: 10.1162/089976601750541778.

Matthew Hausknecht and Peter Stone. Deep recurrent Q-learning for partially observable MDPs. In AAAI Fall Symposium on Sequential Decision Making for Intelligent Agents, 2015.

James B. Heald, Máté Lengyel and Daniel M. Wolpert. Contextual inference underlies the learning of sensorimotor repertoires. Nature, 600(7889):489–493, 2021. doi: 10.1038/s41586-021-04129-3.

James B. Heald, Máté Lengyel and Daniel M. Wolpert. Contextual inference in learning and memory. Trends in Cognitive Sciences, 27(1):43–64, 2023. doi: 10.1016/j.tics.2022.10.004.

Russell Jeter, Yaroslav Molkov, and Dmitrii Todorov. Optimal practice schedules in a dual-rate model of motor adaptation, June 2026. URL 10.5281/zenodo.20721359.

Wilsaan M Joiner and Maurice A Smith. Long-term retention explained by a model of short-term learning in the adaptive control of reaching. Journal of Neurophysiology, 100(5):2948–2955, 2008. doi: 10.1152/jn.90706.2008.

Leslie Pack Kaelbling, Michael L. Littman, and Anthony R. Cassandra. Planning and acting in partially observable stochastic domains. Artificial Intelligence, 101(1–2):99–134, 1998. doi: 10.1016/S0004-3702(98)00023-X.

John W. Krakauer, Alkis M. Hadjiosif, Jing Xu, Aaron L. Wong, and Adrian M. Haith. Motor learning. Comprehensive Physiology, 9(2):613–663, 2019. doi: 10.1002/cphy.c170043.

Jeong-Yoon Lee and Nicolas Schweighofer. Dual adaptation supports a parallel architecture of motor memory. Journal of Neuroscience, 29(33):10396–10404, 2009. doi: 10.1523/JNEUROSCI.1294-09.2009.

Jeong Yoon Lee, Youngmin Oh, Sung Shin Kim, Robert A. Scheidt, and Nicolas Schweighofer. Optimal schedules in multitask motor learning. Neural Computation, 28(4):667–685, 2016. doi: 10.1162/NECO_a_00823.

Tatsuya Nakata and Yuichi Suzuki. Mixing grammar exercises facilitates long-term retention: Effects of blocking, interleaving, and increasing practice. Journal of Educational Psychology, 111 (7):1172–1188, 2019. doi: 10.1037/edu0000336.

Sanmit Narvekar, Bei Peng, Matteo Leonetti, Jivko Sinapov, Matthew E. Taylor, and Peter Stone. Curriculum learning for reinforcement learning domains: A framework and survey. Journal of Machine Learning Research, 21(181):1–50,2020.

Philip I. Pavlik Jr and John R. Anderson. Using a model to compute the optimal schedule of practice. Journal of Experimental Psychology: Applied, 14(2):101–117, 2008. doi: 10.1037/1076-898X.14.2.101.

Tomi Peltola, Mustafa Mert çelikok, Pedram Daee, and Samuel Kaski. Machine teaching of active sequential learners. In Advances in Neural Information Processing Systems 32, 2019.

Siddharth Reddy, Sergey Levine, and Anca Dragan. Accelerating human learning with deep reinforcement learning. In NIPS Workshop on Teaching Machines, Robots, and Humans, 2017.

Doug Rohrer and Kelli Taylor. The shuffling of mathematics practice problems boosts learning. Instructional Science, 35(6):481–498, 2007. doi: 10.1007/s11251-007-9015-8.

Richard A. Schmidt and Robert A. Bjork. New conceptualizations of practice: Common principles in three paradigms suggest new concepts for training. Psychological Science, 3(4):207–217, 1992. doi: 10.1111/j.1467-9280.1992.tb00029.x.

John Schulman, Filip Wolski, Prafulla Dhariwal, Alec Radford, and Oleg Klimov. Proximal policy optimization algorithms. arXiv preprint arXiv:1707.06347, 2017.

Burr Settles and Brendan Meeder. A trainable spaced repetition model for language learning. In Proceedings of the 54th Annual Meeting of the Association for Computational Linguistics, pages 1848–1858, 2016. doi: 10.18653/v1/P16-1174.

John B. Shea and Robyn L. Morgan. Contextual interference effects on the acquisition, retention, and transfer of a motor skill. Journal of Experimental Psychology: Human Learning and Memory, 5(2):179–187, 1979. doi: 10.1037/0278-7393.5.2.179.

Maurice A. Smith, Ali Ghazizadeh, and Reza Shadmehr. Interacting adaptive processes with different timescales underlie short-term motor learning. PLoS Biology, 4(6):e179, 2006. doi: 10.1371/journal.pbio.0040179.

Ronald J. Williams. Simple statistical gradient-following algorithms for connectionist reinforcement learning. Machine Learning, 8(3-4):229–256, 1992. doi: 10.1007/BF00992696.

Laurence A. Wolsey. Integer Programming. John Wiley & Sons, New York, 1998.

Gabriele Wulf and Charles H. Shea. Principles derived from the study of simple skills do not generalize to complex skill learning. Psychonomic Bulletin & Review, 9(2):185–211, 2002. doi: 10.3758/BF03196276.

Xiaojin Zhu. Machine teaching: An inverse problem to machine learning and an approach toward optimal education. In Proceedings of the AAAI Conference on Artificial Intelligence, volume 29, 2015.

